# Validation of ten federated learning strategies for multi-contrast image-to-image MRI data synthesis from heterogeneous sources

**DOI:** 10.1101/2025.02.09.637305

**Authors:** Jan Fiszer, Dominika Ciupek, Maciej Malawski, Tomasz Pieciak

## Abstract

Deep learning (DL)-based image synthesis has recently gained enormous interest in medical imaging, allowing for generating multi-contrast data and therefore, the recovery of missing samples from interrupted or artefact-distorted acquisitions. However, the accuracy of DL models heavily relies on the representativeness of the training datasets naturally characterized by their distributions, experimental setups or preprocessing schemes. These complicate generalizing DL models across multi-site heterogeneous data sets while maintaining the confidentiality of the data. One of the possible solutions is to employ federated learning (FL), which enables the collaborative training of a DL model in a decentralized manner, demanding the involved sites to share only the characteristics of the models without transferring their sensitive medical data. The paper presents a DL-based magnetic resonance (MR) data translation in a FL way. We introduce a new aggregation strategy called FedBAdam that couples two state-of-the-art methods with complementary strengths by incorporating momentum in the aggregation scheme and skipping the batch normalization layers. The work comprehensively validates 10 FL-based strategies for an image-to-image multi-contrast MR translation, considering healthy and tumorous brain scans from five different institutions. Our study has revealed that the FedBAdam shows superior results in terms of mean squared error and structural similarity index over personalized methods, like the FedMRI, and standard FL-based aggregation techniques, such as the FedAvg or FedProx, considering multi-site multi-vendor heterogeneous environment. The FedBAdam has prevented the overfitting of the model and gradually reached the optimal model parameters, exhibiting no oscillations.

## 1. Introduction

Magnetic resonance imaging (MRI) is a non-invasive technique that provides detailed images of the medium without using harmful radiation. This quality makes MRI one of the most popular and versatile imaging techniques in clinical practice used nowadays. In particular, multimodal MRI that incorporates different contrasts is crucial for a comprehensive assessment of the anatomy and function of the human body. The unique information provided by different MRI-based contrasts can enhance the diagnosis in oncology (Zhang et al., 2017), neuropsychiatric disorders like schizophrenia (Salvador et al., 2019), neurodegenerative diseases such as Alzheimer’s (Weiler et al., 2015) and Parkinson’s disease (Bowman et al., 2016) or brain aging (Cherubini et al., 2016). Multiple-contrast MRI can also benefit other MRI-based procedures like brain tumour segmentation (Sun et al., 2019; Zhu et al., 2023) or survival prediction (Sun et al., 2019; Moya-Sáez et al., 2022).

However, obtaining a complete multimodal dataset can be challenging in clinical scenarios due to several factors. The process is often expensive and time-consuming (Zhou et al., 2020), which may make it impractical for imaging patients heavily affected by a disease or those with claustrophobia. As a consequence, scanning defeats can occur, resulting in poor image quality or unfinished acquisitions due to the interruption of the examination (Dai et al., 2020; Yang et al., 2020; Moshe et al., 2024). This further leads to incomplete datasets that negatively impact diagnosis and treatment. Most existing methods do not effectively handle missing modalities, often leading to failures in these situations (Zhou et al., 2020; Moshe et al., 2024).

A possible solution to handle incomplete acquisitions involves medical image synthesis or, specifically, multi-contrast image translation. This process enables the artificial generation of missing modalities without additional scanning procedures. Image synthesis has been successfully applied in clinical settings, particularly in neuro-oncology to improve the survival prediction (Moya-Sáez et al., 2022), segmentation (Yang et al., 2020; Conte et al., 2021) and classification of brain tumours (Moshe et al., 2024), and radiotherapy planning (Brou Boni et al., 2021). Currently, the most widely used approach for image translation involves image-to-image neural networks (Dai et al., 2020; Armanious et al., 2020; Yang et al., 2020; Zhou et al., 2020; Conte et al., 2021; Özbey et al., 2023; Moshe et al., 2024).

The accuracy of deep learning (DL) models heavily relies on the representativeness of the training datasets (Rieke et al., 2020; Aouedi et al., 2023; Aja-Fernández et al., 2023). However, neural networks are frequently trained and evaluated on data selected from a single institution, often using the same data attributes. Consequently, if an organization lacks data on specific pathologies, the trained model may become biased and demonstrate low generalization capabilities when applied to other datasets. Additionally, the generalization ability can be further compromised by variations in data among different centres, which arise from using various devices and acquisition protocols. To address this issue, several initiatives have been launched to systematically gather data from diverse sources to create large and multifarious datasets covering a wide range of diseases and ethnic groups (Sudlow et al., 2015; Wang et al., 2017; Cuggia et al., 2019). Despite these efforts, several challenges remain, especially regarding privacy and the General Data Protection Regulation (Rieke et al., 2020; Darzidehkalani et al., 2022; Aouedi et al., 2023). The solution to this problem may lie in the federated learning (FL) concept (McMahan et al., 2017; Riviera et al., 2023), a decentralized approach to training DL models. This method enables multiple institutions to collaboratively train a neural network model without sharing their sensitive medical data. Instead, each participant conducts training locally and only shares specific characteristics, such as parameters or gradients of the updated models, with a central server. The server then aggregates these characteristics to generate a single global model. Numerous recent studies have demonstrated the potential of FL for various medical imaging tasks, including segmentation (Jiang et al., 2023), reconstruction (Feng et al., 2023), and detection (Bercea et al., 2022). These studies show that a model trained in a decentralized manner achieves significantly better generalization compared to models trained on single datasets.

The FL-based DL has recently been successfully applied to MRI data synthesis, including a generative adversarial model (GAN) that aggregates only the generator parameters while keeping the discriminator parameters locally (Dalmaz et al., 2022), This technique has been recently extended by splitting the generator into a mapper and a downstream synthesizer, shared with a central server, and a local upstream synthesizer (Dalmaz et al., 2024). Another solution with similar assumptions utilizes self-supervised learning, allowing to work with unpaired cross-modality data while effectively handling heterogeneous datasets (Wang et al., 2023). Cycle-Consistent GAN was also utilized for medical image translation within the FL framework (Bdair et al., 2024). Combined with the spatial attention mechanism, it achieved significantly better results compared to traditional convolutional networks like U-Net. However, the existing literature on MRI translation mainly focuses on using more advanced DL models with or without introducing the FL context. Unfortunately, it lacks a comprehensive comparison with existing approaches using multi-sources vendor diverse datasets. This leaves important questions unanswered, such as how to effectively aggregate neural networks trained by different institutions under a heterogeneous environment and whether all involved sites should be treated equally during the training process. Our paper aims to fill this gap and thoroughly compares 10 FL techniques, including three specifically designed for MRI data, focusing on synthesizing structural MRI images. Furthermore, we introduce the FedBAdam, a novel approach to developing FL schemes that integrates two different FL methods. Our analyses employ data from healthy individuals and tumour patients at various stages, particularly glioma patients, collected from multiple institutions and acquired under different acquisition conditions to ensure a representative and accurate reflection of real-world scenarios. The contributions of the paper can be summarized as follows:

1. We introduce the approach of coupling two FL aggregation methods having complementary strengths, namely the FedBN and FedAdam, we obtained a new one called FedBAdam, which significantly outperforms the other methods applied individually to the data translation task. To our best knowledge, this approach has not been tested before and may indicate a new direction for developing FL strategies.
2. We comprehensively validate 10 FL-based strategies for handling image-to-image multi-contrast MRI data translation (i.e., translation between all combinations of T1-weighted, T2-weighted and FLAIR). This includes six foundational FL methods commonly used in medical imaging studies, three techniques specifically designed for MRI data, and one approach we introduce.
3. Engaging six datasets from five different institutions, we simulate FL-based compositions closely resembling real-world conditions. These datasets vary in several ways: they include healthy patients and those with brain glioma under various stages, differ in the number of samples, and have been acquired using different vendors and acquisition protocols.

## 2. Materials and methods

In this research, we use five publicly-available datasets covering T1-, T2-weighted and FLAIR MRI acquisitions:

1. **Human Connectome Project (HCP) WU-Minn** (Van Essen et al., 2013): 104 healthy subjects (35F/69M), aged 22-35y. The subjects were scanned using a customised Siemens 3T Connectome Skyra scanner (Siemens, Erlangen, Germany) equipped with a 32-channel head coil. The dataset includes T1- and T2-weighted scans for each subject acquired under a spatial resolution of 0.7×0.7×0.7 mm^3^.
2. **HCP MGH** (Fan et al., 2015): 26 healthy subjects (13F/13M), aged 20-59y. The subjects were scanned using a Siemens Skyra 3T scanner equipped with a customised 64-channel head coil. The dataset includes T1- and T2-weighted scans acquired using 3D MPRAGE and 3D T2-SPACE protocols, respectively. Other acquisition parameters are (T1-/T2-weighted): accelerated parallel acquisition using GRAPPA protocol under the acceleration factor 2, voxel resolution: 1×1×1 mm^3^/0.7×0.7×0.7 mm^3^.
3. **OASIS-3** (LaMontagne et al., 2019): 80 healthy and senile dementia subjects (49F/31M), aged 42-95y. The subjects were scanned using three Siemens scanners: Vision 1.5T equipped with a 16-channel head coil, TIM Trio 3T and BioGraph mMR PET-MR 3T scanners both using a 20-channel head coil. The dataset includes T1-, T2-weighted and FLAIR scans.
4. **UCSF (UCSF-PDGM-v3)** (Calabrese et al., 2022): 150 subjects (63F/87M), aged 57±15, with histopathologically confirmed gliomas according to WHO classification (grades 2-4). All subjects were scanned using a 3T Discovery 750 GE scanner (GE, Waukesha, WI) with a dedicated eight-channel head coil. The dataset includes T1-/T2-weighted/FLAIR scans acquired using 3D IR-SPGR/3D FSE/3D FSE protocols, respectively. Other acquisition parameters were: voxel resolution: 1×1×1 mm^3^ for T1-weighted scans and 1.2×1.2×1.2 mm^3^ for T2-weighted/FLAIR.
5. **BraTS** (Menze et al., 2015; Bakas et al., 2017; Bakas et al., 2018): further divided into low and high-grade gliomas (LGG, HGG) of 76 and 125 subjects respectively. The dataset covers T1-weighted (sagittal or axial acquisitions, variable slice thickness 1-5mm) T2-weighted (axial acquisitions, variable slice thickness 2-4mm), and FLAIR acquisitions (axial, coronal or sagittal acquisitions).

**Supplementary Table 1** summarises the acquisition parameters used to obtain the aforementioned datasets.

### 2.2. Data preprocessing and postprocessing

The datasets used in the study have been shared by the institutions mostly in a preprocessed form. Below, we summarize the preprocessing and postprocessing steps applied to the data:

1. **HCP WuMinn:** registration of T2- to T1-weighted volumes using a customised FLIRT BBR algorithm, and bias field corrections (Rilling et al., 2012).
2. **HCP MGH:** de-facing and de-earing using a face mask generated with the Freesurfer and an ear mask drawn manually.
3. **UCSF:** registration and resampling of the data to the isotropic 1 mm^3^ resolution 3D space defined by the FLAIR volumes using automated nonlinear registration (Advanced Normalisation Tools; Avants et al., 2009), and skull-stripping (Calabrese et al., 2020), followed by a tumour areas segmentation.
4. **BraTS:** re-orientation to the left-posterior-superior coordinate system, co-registration to a common anatomical template SRI (Rohlfing et al., 2010), resampling to isotropic 1 mm^3^ voxel resolution, skull-stripping, and manual segmentation of tumour tissue.
5. The **OASIS-3** dataset has not been preprocessed.

On top of these, we applied some additional data processing, including:

1. **HCP MGH**: Skull-stripping using the `bet` tool from FSL 6.0.6.4 (Analysis Group, FMRIB, Oxford, UK.; Smith et al., 2004) and linear registration of T2-weighted volumes to T1-weighted space for each subject using the FSL `flirt` with 7 degrees of freedom and then improved the mapping using a nonlinear registration *via* the FSL `fnirt` (Jenkinson & Smith, 2001; Jenkinson et al., 2002).
2. **OASIS-3**: Skull-stripping *via* the FSL `bet` and the registration of T1- and T2-weighted volumes to FLAIR space for each subject with the same procedure as used for the HCP MGH.

The datasets that include only T1-weighted and T2-weighted data (i.e., HCP WU-Minn and HCP-MGH) were transformed into pairs of T1- and T2-weighted, while the datasets also containing FLAIR data (i.e., UCSF, BraTS, and OASIS-3) into triplets of corresponding slices. The voxel values of the images were additionally normalised using the min-max procedure to the range between 0 and 1. Only slices containing a substantial portion of the brain regions were selected for further analysis.

Our experimental scenario reflects a heterogeneous environment due to different acquisition protocols, preprocessing/postprocessing procedures and differences in the number of samples used for training. Moreover, the OASIS-3 dataset is low-quality data, making it a potential malicious client – a common problem in FL (Li et al., 2020a). The capability of an aggregation method to remain robust against the influence of such malicious clients might be an essential factor in choosing the appropriate FL-based approach.

### 2.3 Deep learning-based multi-contrast MRI data synthesis

The U-Net model (Ronneberger et al., 2015; Osman et al., 2022) has been employed to translate MRI data in an image-to-image scenario considering all possible combinations between T1-, T2-weighted and FLAIR acquisitions (see **Supplementary Fig. 1** for details). The encoder starts with an initial block of 64 channels, with the number of channels doubling consistently until reaching 1024 channels at the bottleneck. The decoder mirrors this structure, starting at the bottleneck and producing output images with the original resolution but in a different contrast. We defined the loss function to cover the sum of mean squared error (MSE) and the mean structural similarity index (MSSIM)

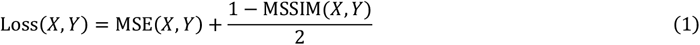

with MSSIM being the mean value of SSIM over the whole training batch including background areas with the SSIM given by (Wang et al., 2004)

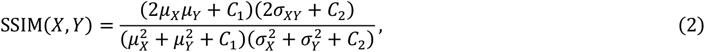

where *μ*_*X*_ and *μ*_*Y*_ are first-order sample (local) moments of the batches *X* and *Y, σ*_*X*_ and *σ*_*Y*_ are the local standard deviations and *σ*_*XY*_ is the local correlation coefficient between the batches *X* and *Y*. The constants in Eq. (2) have been fixed to *C*_1_ = 10^−4^ and *C*_2_ = 9 × 10^−4^.

We use the Adam optimizer (Kingma & Ba, 2014) with the learning rate of 0.001 and the batch size of 32 selected empirically. For each single dataset, the model was trained for 50 epochs, enough for convergence across all translation directions. During the evaluation, the models from the epoch with the lowest validation loss values were utilised. The models trained on a single dataset are referred to as single-trained models (i.e., trained in a non-FL way). The datasets were divided in the following proportions for training, testing and validation: 75%, 15%, 10%.

### 2.4 Deep learning model aggregation methods in a federated learning way

Ten different aggregation techniques have been implemented and compared, all under the same experimental setups: 4 local epochs and 32 global rounds (training iterations). The fraction fit was set to 1.0, meaning all clients were included in every training iteration. The aggregation methods considered in the study are as follows (see section 3 in the **Supplementary materials** for technical details):

1. **FedAvg** (McMahan et al., 2017): a weighted average of the model parameters;
2. **FedMean**: an arithmetic mean (not-weighted average);
3. **FedProx** (Li et al., 2020b): designed to deal with the heterogeneity of computing devices; incorporates *L2* regularisation in the loss function computed relative to the global model weights, and randomly sampled stragglers (a subset of clients trained for different number of epochs);
4. **FedAdam, FedAdagrad** (Reddi et al., 2020): adaptive optimization methods;
5. **FedCostWAvg** (Mächler et al., 2021): includes the loss change between two consecutive rounds as the weight during the average computation;
6. **FedPIDAvg** (Mächler et al., 2023): an iterative improvement of FedCostWAvg, which additionally considers the loss from the last five rounds;
7. **FedBN** (Li et al., 2021): weighted average of the model parameters as the FedAvg, but excludes normalisation layers;
8. **FedMRI** (Feng et al., 2023): model is divided into the global encoder and local decoders;
9. **FedBAdam**: a mixture of FedAdam and FedBN methods, i.e., includes momentum in the aggregation scheme as the FedAdam, but skips the batch normalisation layers identically to FedBN.

The methods yielding a global model are henceforth referred to as global methods (i.e., FedAvg, FedMean, FedProx, FedAdam, FedAdagrad, FedCostWAvg and FedPIDAvg). The personalised methods FedBN, FedMRI, and FedBAdam result in personalised models for each client, potentially enhancing the model performance in heterogeneous data environments. However, these methods lack a global model, thus the evaluations were performed on clients’ models. For all methods considered in the study, the final model (i.e., the model from the last global round) has been selected, facilitating a fair comparison. The methods have been implemented using the Flower 1.4.0 framework (Beutel et al., 2020).

### 2.5 Quality metrics

For evaluation purposes, the MSSIM metric and the MSE are two indicators of model output quality. Both metrics are calculated exclusively over the brain mask, including the white matter, gray matter and the cerebrospinal fluid. Considering the MSSIM computed only over the brain structure prevents artificial score boosting (score inflation) that appears due to accurate background prediction (see Fig. 3 in Liu et al., 2021). Besides calculating the MSSIM over the brain region, we obtain the MSSIM over a trimmed part of the brain where the tumour mask is located. To his end, five extra data points margin has been included in the bounding box of reference brain tumour masks to preserve the surrounding area of the tumour. This metric emphasises tumour translation between multiple-contrast MRI data – a crucial part of the quality assessment. We compute this metric only for datasets providing subjects with tumours (i.e., BraTS and UCSF).

## 3. Experimental results

### 3.1 Non-FL-based image translation

We start demonstrating our results from non-FL image-to-image translation to highlight the trouble of image translation between clients characterized by different acquisition protocols. In **Figs. 1a** and **2b**, we visually inspect the translation between T1- and T2-weighted MRI data for a representative axial slice in a non-FL way (see **Supplementary Figs. 2** and **3** for translations *to* and *from* FLAIR domain). These figures present the synthesized images in the left panels and the relative errors compared to the reference images in the right panels considering the following scenario. The first row in the left panel presents the input data acquired at different clients, while the second row illustrates the target data from the same corresponding clients (the domain to which the input data has to be translated). Then, each consecutive row starting at position 3 presents the results of image translation for each client (annotated in the column header) considering the training procedure with data annotated in the row header. The diagonals in the right panels of **Figs. 1a** and **1b** demonstrate the best performance for each of the testset has been achieved for the models trained on the samples coming from the same data source. More intense error colours for all the other datasets are the first hint of their poor generalization capabilities between the clients. Indeed, the most destructed visual results are notable for the model trained on the HCP Wu-Minn dataset for T2-weighted→T1-weighted translation direction (see **Fig. 1b**; right panel). This network faced significant challenges when translating images for UCSF and BraTS data sets due to substantial differences in data characteristics, including spatial resolution, noise levels, and pixel intensities for certain tissues.

**Figure 1.**
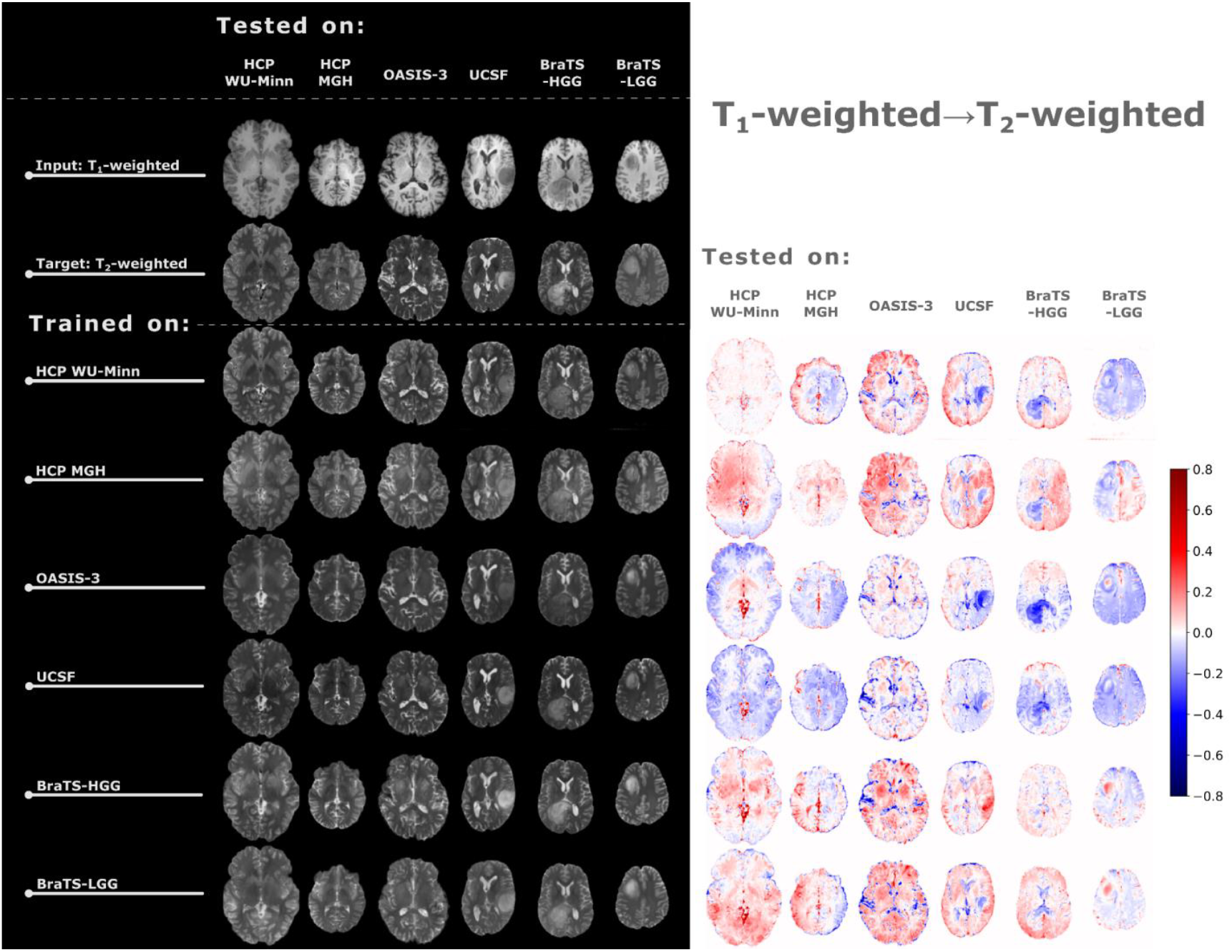
Image translation results between T1- and T2-weighted MRI in a non-FL way (left panel) and the relative errors calculated between the synthesized data and target images. The first row in the left panel presents the input data, and the second row indicates the target domain (the domain to which the input data is translated). Each subsequent row depicts the results of image translation for the clients (each annotated in the column header) given the training procedure conducted with the data set annotated in the row header.

**Figure 1b.**
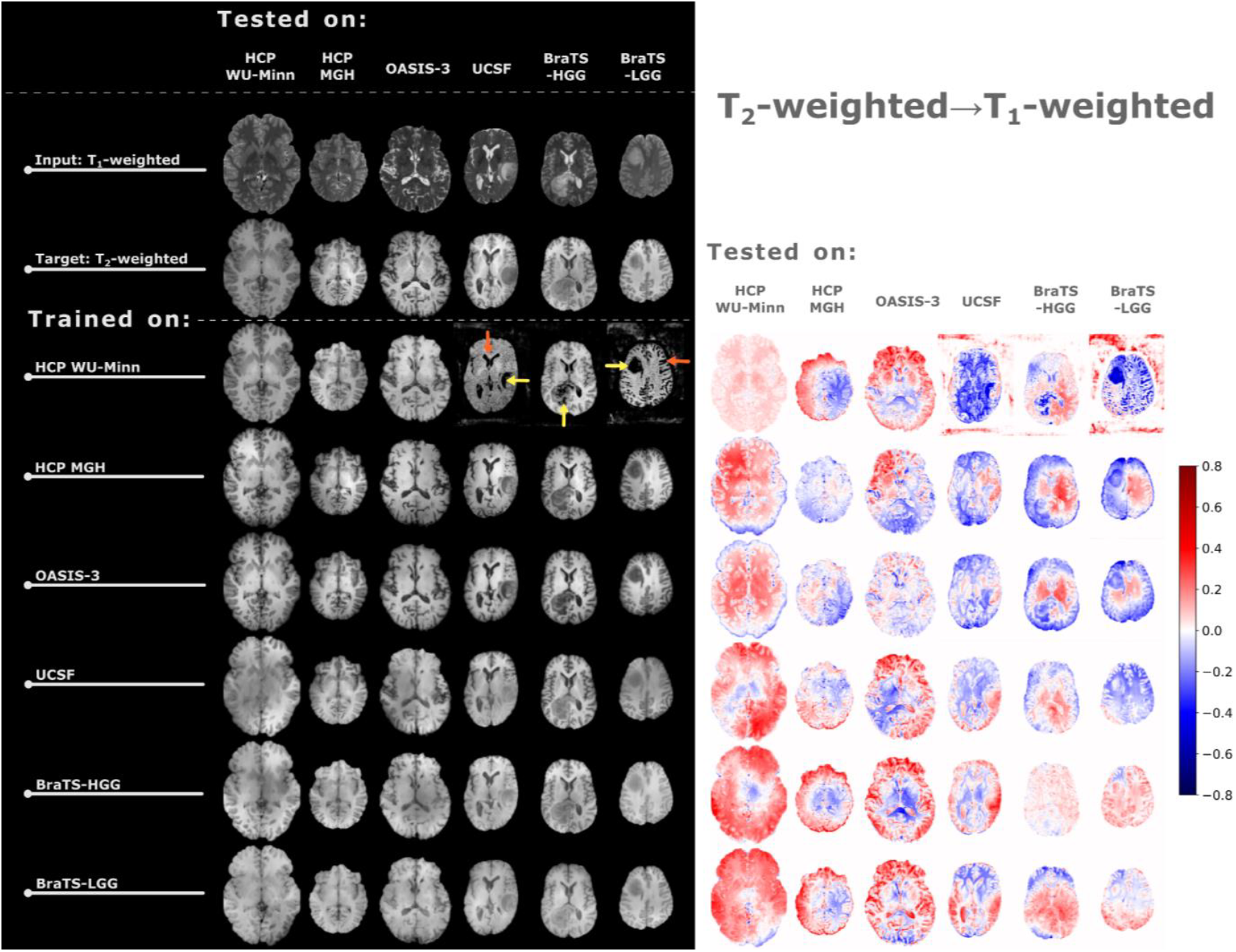
(cont.) Yellow arrows indicate artifacts in the brain tumor area, while orange arrows identify artifacts in healthy tissues and the cerebrospinal fluid.

The neural network struggles by mistakenly identifying some brain regions, especially gliomas, as the background area, by setting the values close to zero, i.e., almost black areas (see row 3 in **Fig. 1b**). The relative errors in translation from T1- and T2-weighted data to the FLAIR domain are even more exposed than translation between T1- and T2-weighted data (see **Supplementary Figs. 2a** and **3a**). This simple experiment shows the need for knowledge generalization in the neural networks due to heterogeneity between the training and testing datasets.

### 3.2 FL-based image translation learning curves

From now on, we handle the image translation task in the clutches of the FL framework considering the methods presented in section 2.4. First, in **Fig. 2**, we illustrate the averaged loss function values defined by **Eq. (1)** as a function of the global round number. We can observe that the choice of aggregation method affects both the convergence rate (training speed) and the model performance, with more sophisticated aggregation methods being capable of achieving lower loss function values. The best results are obtained for the FedBAdam method for each contrast combination, followed by the second-best FedMRI. Among the remaining methods, the FedBN reaches the lowest loss value. Considering the global methods, the lowest loss function is usually obtained with the FedAdam, but the loss differences are minor and vary depending on the translation direction. This divergence is particularly evident with FedProx, which performs the best among the global methods for FLAIR→T2-weighted translation and the worst for T2-weighted→T1-weighted translation. Another important characterization of the aggregation methods revealed by the learning curves is the difference between the adaptive optimization methods (i.e., FedAdam, FedAdagrad). Despite being designed with similar principles, they perform differently–FedAdam is relatively smooth, whereas FedAdagrad is very unstable in the loss values as a function of the global round.

**Figure 2.**
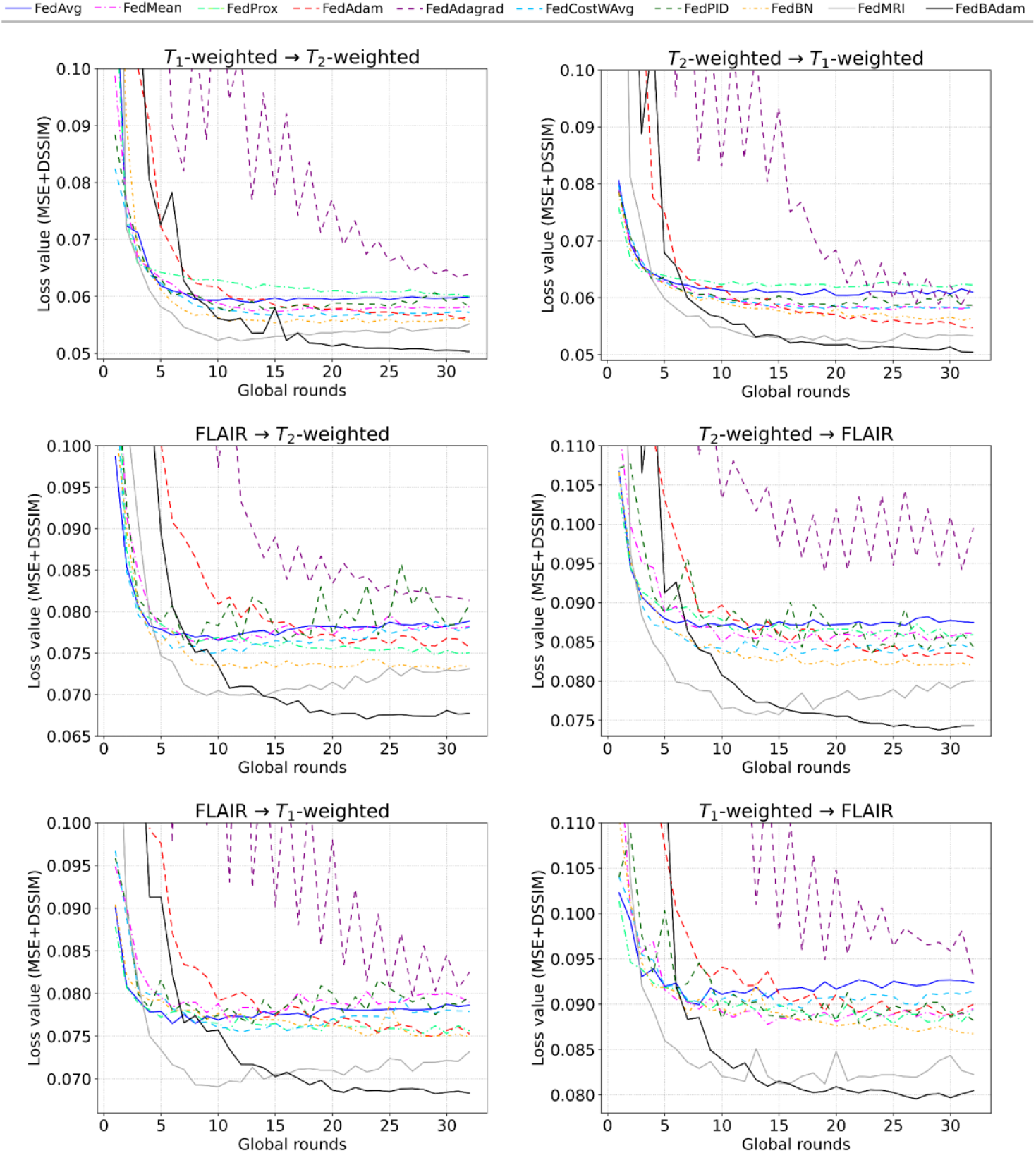
The change of the loss values as a function of the global round number (global epochs) in the FL-based training. Each plot presents an averaged loss function calculated across all six clients involved in the FL-based training. The annotation on top of the plot shows the direction of image translation.

Finally, concerning the convergence speed, FedAdam and FedBAdam gradually reach satisfactory loss levels compared to other methods. On the other hand, FedMRI is characterized by the fastest drop, typically attaining its minimal value within 10 global rounds. However, FedMRI is the most prone to overfitting for all translation directions. This unwanted tendency can also be noticed in some translation directions for FedAvg, FedPID, and FedCostWAvg, though to a lesser extent.

The overfitting issue is further investigated for two personalized methods, namely the FedMRI and FedBAdam, by analyzing the learning curves of each client separately (see **Fig. 3** and **Supplementary Fig. 4**). We can see the FedMRI method overfits only for some clients, most significantly visible for the OASIS-3, to be considered as the malicious client. The OASIS-3 client learning curve is unstable in terms of convergence and starts increasing at some global round for the FedMRI, which is indubitably an undesired behaviour. Contrary, for the FedBAdam approach, all the clients steadily decrease their loss values with the global rounds, though the OASIS-3 is still considered as the worst client.

**Figure 3.**
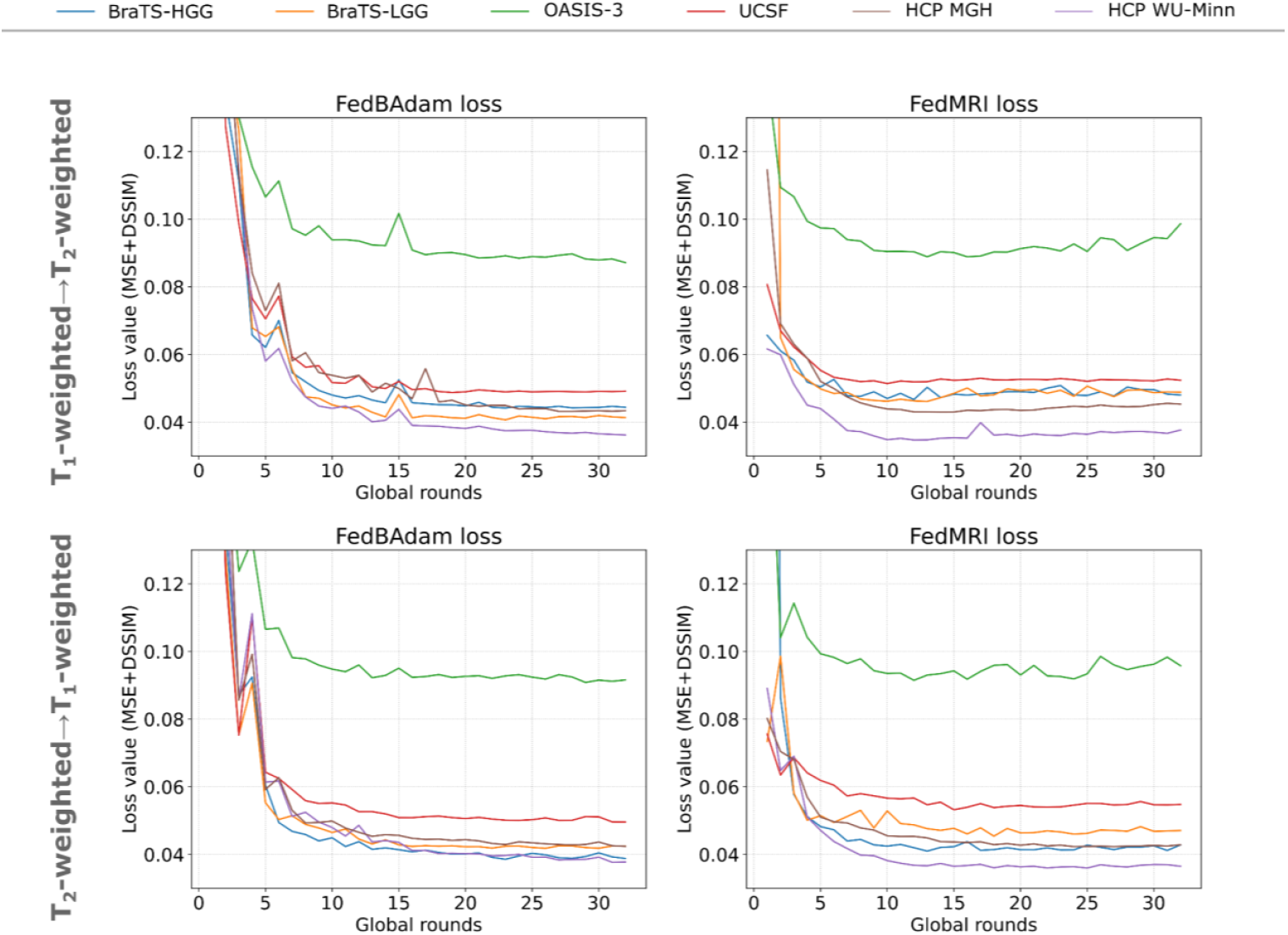
The change of the loss values as a function of the global round number (global epoch) across the clients involved in the FL-based training for two selected personalized methods (i.e., FedBAdam and FedMRI). The annotations on the left show the direction of image translation.

### 3.3 FL-based image translation: qualitative results

In the next experiment, we present visual inspection of image-to-image translation for the network trained in the FL-based way. We involve all clients and train the model using all 10 FL-based approaches previously presented in section 2.4 The results for translation between T1- and T2-weighted data are demonstrated in **Figs. 4a** and **4b**. Additionally, in **Supplementary Figs. 5** and **6**, we illustrate translations *to* and *from* the FLAIR domain.

**Figure 4a.**
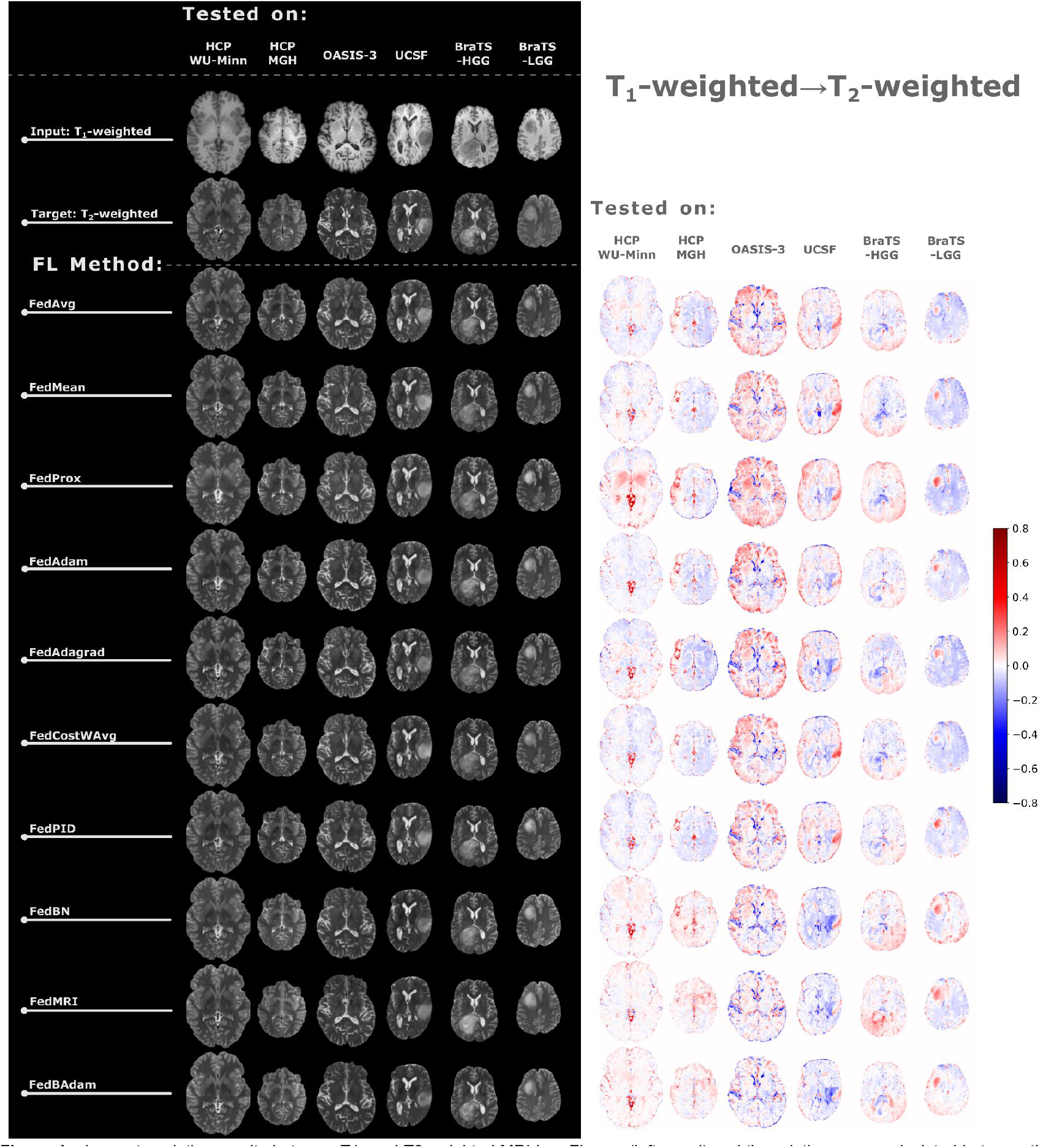
Image translation results between T1- and T2-weighted MRI in a FL way (left panel) and the relative errors calculated between the synthesized data and target images. The first row in the left panel presents the input data, and the second row indicates the target domain (the domain to which the input data is translated). Each subsequent row depicts the results of the image translation task under the FL-based architecture annotated in the row header for the clients annotated in the column header.

**Figure 4b.**
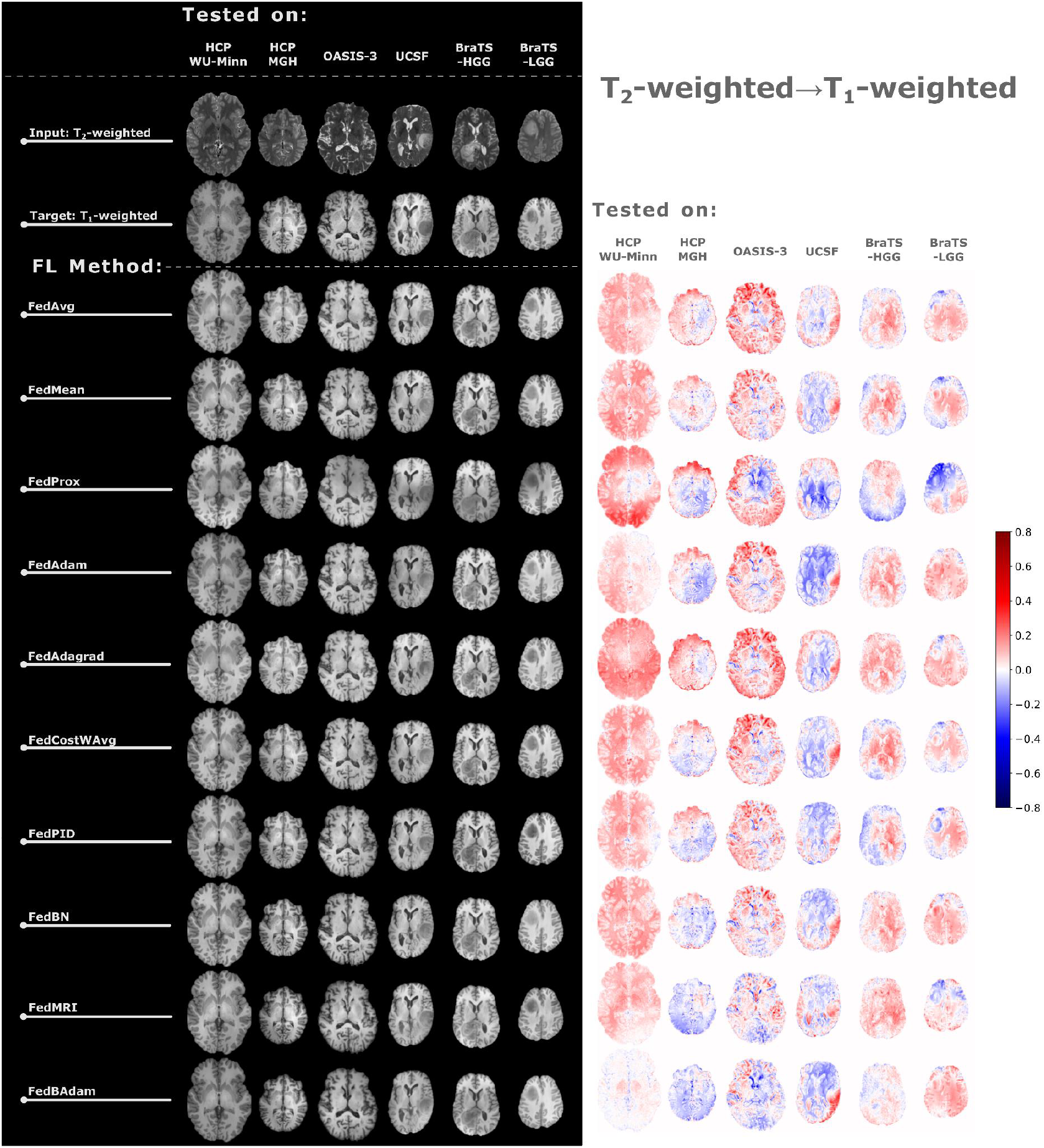
(cont.)

First, translation results in the FL-way are characterized by a smaller relative error (cf. **Fig. 4a** to **Fig. 1a**, for instance). This is a clear example of how neural networks trained traditionally may lack generalization capabilities. On the contrary, the network trained in an FL way can generalize the knowledge from all sources, though, as noted previously, the OASIS-3 client may be considered malicious in the model aggregation procedure. The synthesised slices can be considered satisfying, but they are most inaccurate in the glioma areas, which are particularly important for proper tumour segmentation. The error intensities are the highest for the OASIS-3 dataset, possibly due to a high noise present in the data.

### 3.4 FL-based image translation: quantitative results

To investigate the results further in an image-to-image translation task, we obtain quantitative metrics illustrating the MSE (**Fig. 5**) and MSSIM (**Figs. 6** and **7**) computed for datasets synthesized in non-FL and FL ways. The results presented in **Fig. 1** confirm that the models trained in a non-FL scenario on datasets different than those used for the testing procedure characterize the quality deterioration of synthesized data, which is explained here in terms of increased MSE in non-diagonal cells. Regarding the data synthesized in a FL scenario, consistently, the best method in terms of the MSE computed over the brain areas is the FedBAdam, usually followed in varying order by FedBN, FedMRI and FedAdam. The MSEs present relatively high standard deviation values. Upon closer investigation, it was observed that this phenomenon is caused by outliers (i.e., individual samples significantly deviating from the average) rather than the instability of the translation model.

**Figure 5.**
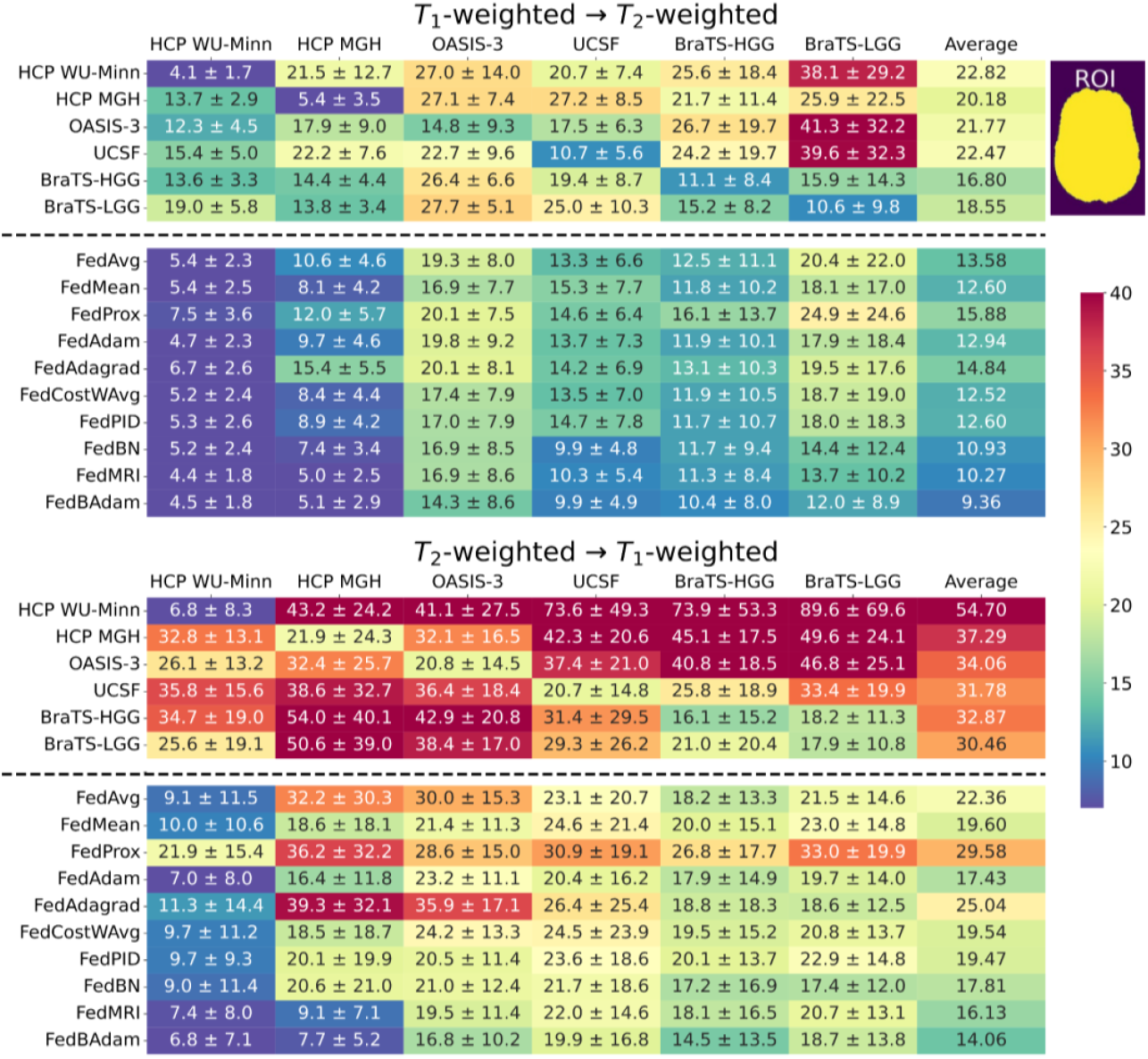
The MSE (multiplied by a factor of 1000) calculated for synthesized data over the brain region in T1-weighted→T2-weighted translation direction (top table) and T2-weighted→T1-weighted translation. In both tables the first six rows cover the translation results in a non-FL way, while the remaining rows represent the ten tested FL methods. Each cell represents the MSE ± standard deviation of the error. The last column presents the average computed across all datasets given the training dataset or FL variant.

**Figure 6.**
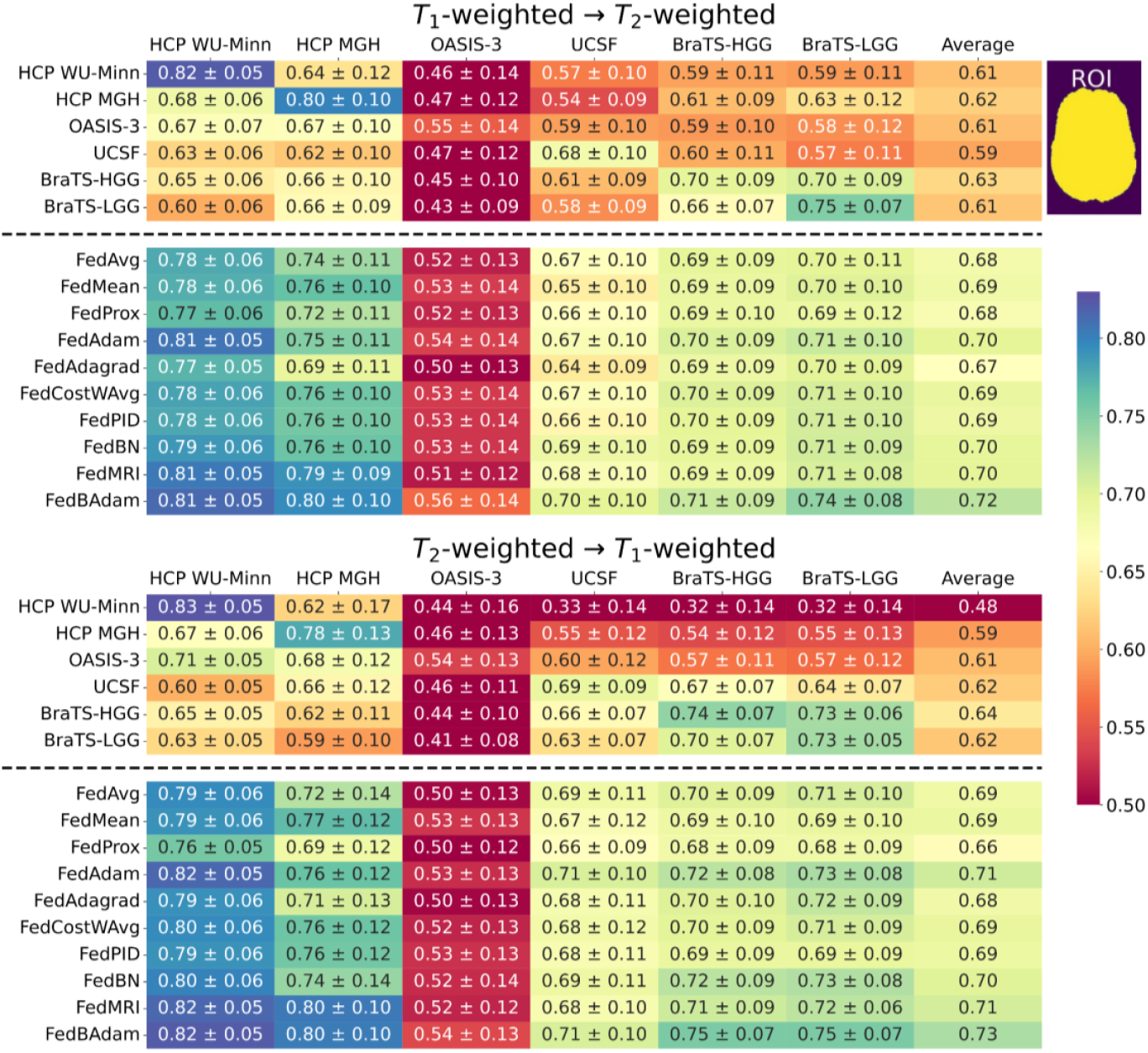
The MSSIM calculated for synthesized data over the brain region in T1-weighted→T2-weighted translation direction (top table) and T2-weighted→T1-weighted translation. In both tables the first six rows cover the translation results in a non-FL way, while the remaining rows represent the ten tested FL methods. Each cell represents the MSSIM ± standard deviation of the SSIM metric. The last column presents the average computed across all datasets given the training dataset or FL variant. The higher the MSSIM the better.

Among the global methods, FedBAdam archives the highest MSSIM values (see **Fig. 6**), slightly surpassing two personalised methods (FedMRI and FedBN). FedAdagrad and FedProx yield the lowest MSSIM for translation between T1-weighted and T2-weighted contrasts, with FedProx’s poor performance mainly caused by a significant drop in the performance for the HCP MGH dataset. Furthermore, FedProx has the highest MSE score for both directions, generally indicating the weakest performance. On the contrary, in the supplementary materials (see **Supplementary Fig. 8**), FedProx emerges to achieve one of the best MSSIM values among global methods for translation with FLAIR. Particularly, superior FedProx performance is highlighted by the tumour-trimmed MSSIM for the T1-weigthed→FLAIR.

All in all, the “average” results for the MSE and MSSIM metrics computed on the brain mask (the last column in **Figs. 5** and **6**) indicate an evident advancement of FL-based over non-FL models. Even the rudimentary FedMean technique outperforms all non-FL models. Compared to the learning curves presented in **Fig. 2**, the performance difference between global and personalised FL methods is less pronounced, with some results being closely aligned.

Concerning the tumour-trimmed MSSIM (see **Fig. 7**), models trained on the data that contain only healthy subjects considerably struggle in the glioma areas. Meanwhile, all FL-based models aggregate information from the other datasets and perform comparably or better regarding the single-trained models with pathological data samples. For T2-weighted→T1-weighted translation, there are noticeable differences between the averaged FL results. Numerous aggregation methods result in a model with worse results than the model trained exclusively on BraTS-HGG, with only FedAdam and FedBAdam indicating superior results. On the other hand, for the opposite translation, all FL and some non-FL-based models perform similarly to BraTS-LGG, with their average scores around MSSIM=0.6. There is an evident performance gap for BraTS-LGG evaluation, where the model trained on BraTS-LGG data achieves MSSIM=0.72, while the closest FL-based result has been obtained for the FedBAdam under the score of MSSIM=0.66. Overall, across most evaluations, FL models perform comparably to the models trained in a non-FL way on their respective datasets.

**Figure 7.**
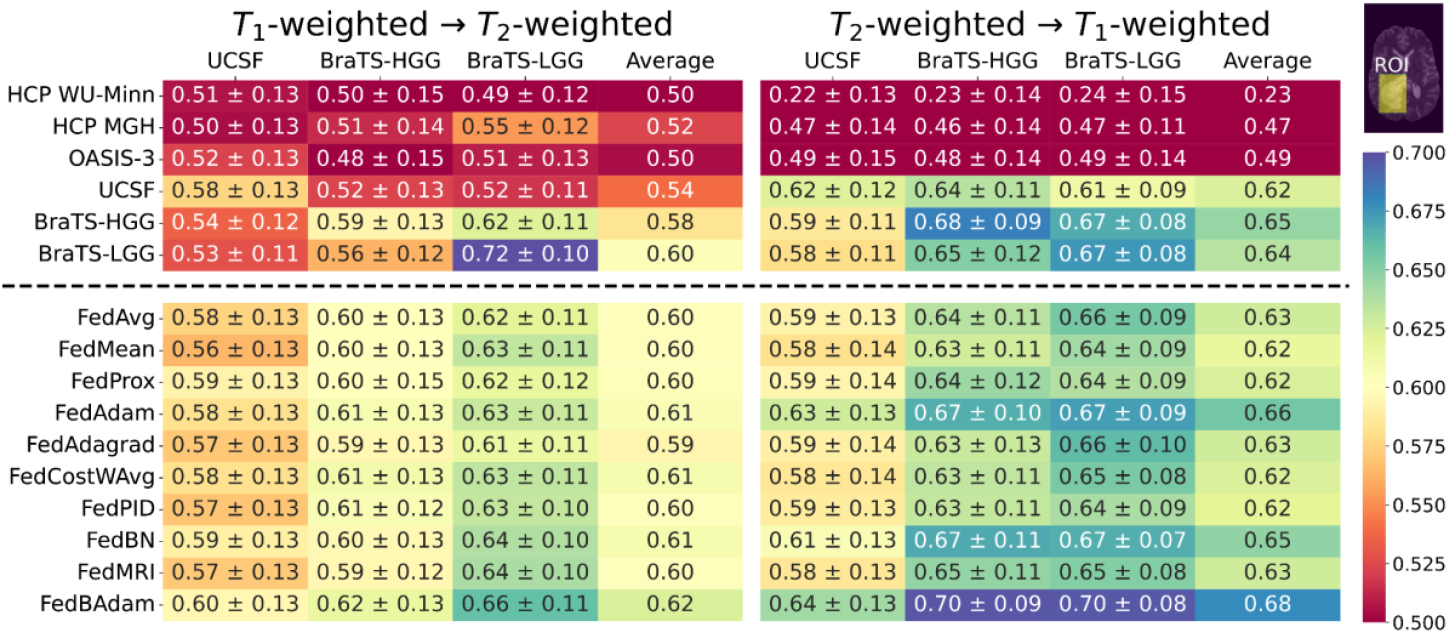
The MSSIM calculated for synthesized data over the tumours areas in T1-weighted→T2-weighted translation direction (top table) and T2-weighted→T1-weighted translation. In both tables the first six rows cover the translation results in a non-FL way, while the remaining rows represent the ten tested FL methods. Each cell represents the MSSIM ± standard deviation of the MSSIM metric. The last column presents the average computed across all datasets given the training dataset or FL variant. The higher the MSSIM the better.

Lastly, but certainly important, some FL-based methods yield better results on specific datasets than models exclusively trained on the same-source data regarding the tumour-trimmed MSSIM metric (cf. the MSSIM in **Fig. 7** computed for the UCSF and BraTS-HGG datasets trained in a non-FL way versus the FL-way). This implies that the federated approach has effectively leveraged information from other datasets, which enhanced the shared model’s performance.

## 4. Discussion

The experimental results presented in the previous section demonstrate that FL-based models can achieve better generalisation capabilities than those trained in a non-FL way (cf. **Fig. 1** and **Fig. 4**). The FL-based training enhances model performance while maintaining data privacy. It not only leads to better overall performance, as illustrated in the “average” column in **Figs. 5** and **6**, but also enables the model to leverage information from the other datasets and, in some cases, even outperform the models trained and tested exclusively on a single dataset (cf. the results in **Fig. 5** for the HCP MGH data trained using the FedMRI and FedBAdam). However, we must make it clear that although “on average” the FL-trained models outperform those trained using a single dataset, one must carefully select the aggregation technique that is appropriate to the datasets’ characteristics across the clients involved in the training (e.g., the FL aggregation technique that handles heterogeneous data correctly).

The aggregation method strongly affects the convergence rate of FL-based trained networks and the final model performance, as observed in the learning curves in **Fig. 2**. Personalised methods (FedBAdam, FedMRI, and FedBN) usually yield the most satisfactory models since they are tailored for each client. Generally, the more sophisticated aggregation method gives better results, with FedAvg performing worse than the other methods most of the time (excluding FedAdagrad). However, these improved methods can increase the complexity of optimising the hyperparameters, such as for FedAdagrad and FedPID. An important characterization of various aggregation methods revealed by the learning curves in **Fig. 2** is the difference between the pair of adaptive optimization methods (FedAdam, FedAdagrad). An important characterization of various aggregation methods revealed by the learning curves in **Fig. 2** is the difference between the pair of adaptive optimization methods (FedAdam, FedAdagrad). Despite both being designed with similar principles, they perform differently—FedAdam is relatively good, whereas FedAdagrad is very unstable. These oscillations are probably due to hyperparameter optimization, which can complicate an appropriate method choice (Zhang et al., 2024) Similar issues occur with FedCostWAvg and FedPIDAvg. Although FedCostWAvg surpasses the baseline method (FedAvg) and sometimes reaches a similar loss level as the best global method (FedAdam), its upgraded version (FedPIDAvg) is unstable for specific translation directions and translates to perform worse than FedAvg (see FLAIR→T2-weighted results in **Fig. 2**).

Among the global methods, selecting the one that is the most accurate for the image-to-image translation task is difficult. Despite changing only the directions of the translation, the performance varies for different aggregation methods, indicating that in the case of real-life deployment, it is advisable to test all candidate FL methods under different hyperparameters. Given the multitude of experiments required, it is essential to analyse the specifics of certain applications. For example, if the goal is rapid convergence, methods having a substantial private network component (like FedMRI; Feng et al., 2023) should be prioritised while being aware of possible overfitting. As illustrated in **Fig. 3**, overfitting can involve merely some clients, therefore, it is advised to analyse the loss changes for each client individually to control this problem rigorously.

In the initial data preparation stage, we used the standard min-max method to normalise individual slices. However, this technique has several limitations (Nyul et al., 1999). Firstly, individual images, even from the same data volume, may differ in the presence or absence of particular tissue types. For example, cerebrospinal fluid may not be visible in some slices, or tumorous tissue may only be present in specific images. Consequently, the maximum value of the MRI signal used for normalisation will also differ, leading to variations in contrast levels within the same dataset. This can pose challenges for training deep learning models and may underestimate their performance for the test set. Secondly, all the datasets we used had undergone prior skull removal, but we cannot guarantee the perfect skull stripping results across all datasets. Consequently, single pixels remaining from the skull area keep outlying MRI signal values, which are also used for min-max normalisation. This further distorted the contrast, even among data from the same subject. In the future, it is necessary to consider alternative methods for normalising the data. Rather than applying this process to individual slices, the entire volumes should undergo normalisation to avoid any distortion caused by the differing maximum values observed within a single subject. Using the value of a selected quantile (e.g., 95th percentile) instead of the maximum value could be an optimal solution, as it would enable filtering outliers, such as those remaining after skull removal. Additionally, it may be beneficial to utilise commonly used techniques for normalising MRI data, such as Z-score, or methods designed explicitly for MRI data (Nyul et al., 1999; Shinohara et al., 2014), which have shown promising results when combined with deep learning approaches (Reinhold et al., 2019; Jacobsen et al., 2019; Alnowami et al., 2022).

Describing a precise goal and metric adaptation is significant to clearly distinguish the best FL methods because incorrectly adopted ones perform comparably to the models trained in a non-FL way. For example, in this study, only FedBAdam achieved consistently the highest scores regarding tumour-trimmed MSSIM (see **Fig. 7**). Despite the optimal performance of the final FL models, some metric values were better in earlier training rounds. Selecting the best model in FL-based training is more complex than classically trained models, especially for personalised methods. The goal is to achieve a well-trained model with robust generalisation capabilities, requiring evaluation across all the datasets, which is a significant computation cost increase and violates privacy concerns. Additionally, utilising the final models tests the method’s vulnerability to overfitting, an essential concern during strategy selection.

In this paper, we employed the U-Net structure as a proof-of-concept due to its simplicity and no risk of generating hallucinations, a critical side-effect particularly important for image translation (Cohen et al., 2018). Despite these reasons, it would be advisable to evaluate FL-based techniques with other deep learning architectures (Yang et al., 2020; Yan et al., 2022; Özbey et al., 2023). Such analyses are particularly important as different types of neural networks, such as GANs, have demonstrated promising results in the FL-based environment (Dalmaz et al., 2022; Wang et al., 2023; Dalmaz et al., 2024; Bdair et al., 2024). Conducting these comparisons would provide a detailed understanding of whether the commonly employed FL approach, where the model is divided into a local component and a part shared with the central server, is truly the most effective. Furthermore, it would be interesting to verify FL-based techniques under various similarity measures (Dohmen et al., 2024) and to explore image translation in an unpaired scenario (Amirkolaee et al., 2022).

Last but not least, the FedBAdam results indicate that combining different methods is reasonable and can lead to better results. In the case of choosing between two well-adapted, distinct methods, using both might be the most profitable solution, as they have complementary strengths. Extracting beneficial properties of currently existing methods and combining them can create an optimal solution for a given task, potentially eliminating the need to invent a dedicated method from scratch. Any new strategy should be compared with a variety of established ones, not only FedAvg or FedProx. The FL field is rapidly evolving, so reliance solely on base approaches as reference points might be no longer advisable.

## 5. Conclusions

This study has successfully integrated deep learning-based image-to-image MRI translation with 10 FL architectures to address data privacy concerns. Depending on the metric used for evaluation, most FL methods either outperformed or achieved similar results compared to models trained on a single-source dataset. Notably, they never strongly malfunctioned, occasionally observed in classically trained models. The FL-based training yields models that handle data heterogeneity more effectively, being a key factor in collaborative training. Reducing the need for perfectly unified training data facilitates broader participation among institutions in federated training processes. Nevertheless, selecting the appropriate aggregation method remains challenging since more advanced techniques may lead to a more accurate model but increase the complexity of method adaptation. As with centralized deep learning depending on the chosen metric, varying methods emerge to be the most promising, highlighting the need for a precise and complete objective function definition. Furthermore, the study indicates that a combination of different approaches could offer the most suitable solution by leveraging the strengths of each method, which, to our best knowledge, has not been tested before. These findings highlight the promise of FL as a methodology with significant prospects for further exploration and development in heterogeneous MRI environments.

## Data availability

Data were provided in part by OASIS, OASIS-3: Longitudinal Multimodal Neuroimaging: Principal Investigators: T. Benzinger, D. Marcus, J. Morris; NIH P30 AG066444, P50 AG00561, P30 NS09857781, P01 AG026276, P01 AG003991, R01 AG043434, UL1 TR000448, R01 EB009352. AV-45 doses were provided by Avid Radiopharmaceuticals, a wholly owned subsidiary of Eli Lilly. Data collection and sharing for this project was provided by the Human Connectome Project (HCP; Principal Investigators: Bruce Rosen, M.D., Ph.D., Arthur W. Toga, Ph.D., Van J. Weeden, MD). HCP funding was provided by the National Institute of Dental and Craniofacial Research (NIDCR), the National Institute of Mental Health (NIMH), and the National Institute of Neurological Disorders and Stroke (NINDS). HCP data are disseminated by the Laboratory of Neuro Imaging at the University of Southern California.

## Acknowledgements

The numerical experiment was possible through computing allocation on the Ares and Athena systems at ACC Cyfronet AGH under the grants PLG/2023/016117 and PLG/2024/016945. This work is supported by the European Union’s Horizon 2020 research and innovation programme under grant agreement No 857533 and the International Research Agendas Programme of the Foundation for Polish Science No MAB PLUS/2019/13. This publication was created within the project of the Minister of Science and Higher Education “Support for the activity of Centers of Excellence established in Poland under Horizon 2020” on the basis of the contract number MEiN/2023/DIR/3796. Tomasz Pieciak acknowledges the Polish National Agency for Academic Exchange for grant PPN/BEK/2019/1/00421 under the Bekker programme and the Ministry of Science and Higher Education (Poland) under the scholarship for outstanding young scientists (692/STY/13/2018).

