## Supplementary materials for "Validation of ten federated learning strategies for multi-contrast image-to-image MRI data synthesis from heterogeneous sources"


### 1. Datasets

**Supplementary Table 1.** The summary of experimental parameters used to acquire the datasets considered in the study.

| | T1-weighted | | | T2-weighted | | | FLAIR | | |
|---|---|---|---|---|---|---|---|---|---|
| **Dataset** | **Voxel size [mm³]** | **Field of view [mm²]** | **Acquisition matrix** | **Voxel size [mm³]** | **Field of view [mm²]** | **Acquisition matrix** | **Voxel size [mm³]** | **Field of view [mm²]** | **Acquisition matrix** |
| **HCP WU-Minn** | 0.7×0.7×0.7 | 224×224 | 320×320 | 0.7×0.7×0.7 | 224×224 | 320×320 | N/A | | |
| **HCP MGH** | | 256×256 | 256×256 | 0.7×0.7×0.7 | 224×224 | 320×320 | N/A | | |
| **OASIS-3** | 1×1×1 | 256×256 | 256×256 | N/A | N/A | N/A | N/A | N/A | N/A |
| **UCSF** | 1×1×1 | 256×256 | 256×256 | 1×1×1.2 | 256×256 | 256×256 | 1×1×1.2 | 256×256 | 256×256 |
| **BraTS** | slice thickness: 1-5mm | N/A | N/A | slice thickness: 2-4mm | N/A | N/A | N/A | N/A | N/A |

### 2. U-Net model used for image-to-image translation

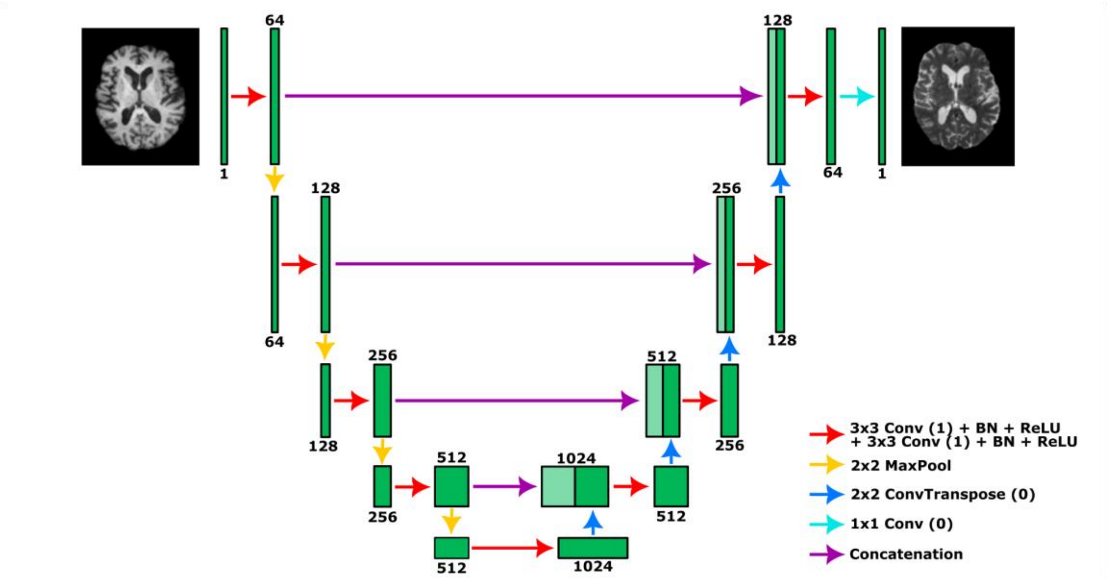


**Supplementary Figure 1.** The schema of the convolution neural network (U-Net) used for image-to-image translation models.

### 3. Federated learning algorithms used in the experiments

#### 1.1 FedAvg (McMahan et al., 2017):

The global model parameters are calculated as follows:

$$\boldsymbol{\Theta}^{r+1} = \sum_{c=1}^{C} w_c \boldsymbol{\Theta}_c^r, \tag{1}$$

where $\boldsymbol{\Theta}_c^r$ denotes the parameters of client $c$ local model at federated round $r$ and $w_c$ is the aggregation weight, defined as $w_c = n_c/N$ with $n_c$ being the number of examples at each client $c$, and $N$ the total number of examples, $N = \sum_{c=1}^{C} n_c$.

#### 1.2 FedMean:

The global model parameters are determined similarly to the FedAvg method, but without using a weighting factor:

$$\boldsymbol{\Theta}^{r+1} = \frac{1}{C}\sum_{c=1}^{C} \boldsymbol{\Theta}_c^r. \tag{2}$$

#### 1.3 FedProx (Li et al., 2020b):

The aggregation process follows the same approach as the FedAvg algorithm, but each client c optimizes a regularized loss function that includes a proximal term $\mu$:

$$\min_{\boldsymbol{\Theta}_c^r} \mathcal{H}_c(\boldsymbol{\Theta}_c^r, \boldsymbol{\Theta}^r; \mathcal{D}_c) = \mathcal{L}_c(\boldsymbol{\Theta}_c^r; \mathcal{D}_c) + \frac{\mu}{2}\|\boldsymbol{\Theta}_c^r - \boldsymbol{\Theta}^r\|^2, \tag{3}$$

where $\boldsymbol{\Theta}^r$ denotes the parameters of global model at federated round $r$, $\mathcal{L}_c$ a local loss function, and $\mathcal{D}_c$ is the private dataset. The variant implemented in this study causes stragglers to train a variable number of local epochs – inversely proportional to the number of training samples. From the FedProx parameters suggested in the original article ($\mu$={0.001, 0.01, 0.1, 1} and stragglers={0, 0.5}), the best results were achieved for: $\mu$=0.001, stragglers=0.5.

#### 1.4 FedAdam, FedAdagrad (Reddi et al., 2020):

The global model parameters are updated as follows:

$$\boldsymbol{\Theta}^{r+1} = \boldsymbol{\Theta}^r + \eta \frac{\boldsymbol{m}^r}{\sqrt{\boldsymbol{v}^r} + \tau}, \tag{4}$$

where $\tau$ controls the algorithm's degree of adaptivity, $\eta$ is the learning rate, and $\boldsymbol{m}^r$ is calculated as:

$$\boldsymbol{m}^r = \beta_1 \boldsymbol{m}^{r-1} + (1-\beta_1)\Delta^r \tag{5}$$

with $\Delta^r = \frac{1}{C}\sum_{c=1}^{C}(\boldsymbol{\Theta}_c^r - \boldsymbol{\Theta}^r)$, and $\beta_1$ being a decay parameter, $\beta_1 \in [0,1)$. The parameters used in the study follows the default configuration setup from the Flower framework except for $\tau$, which was increased for the aggregation stability (Beutel et al., 2020): $\tau$ = 0.001, $\eta$ = 0.1, $\eta_c$ = 0.1, FedAdam: $\beta_1$ = 0.9, $\beta_2$ = 0.99.

For **FedAdam** $\boldsymbol{v}^r$ is determined as:

$$\boldsymbol{v}^r = \beta_2 \boldsymbol{v}^{r-1} + (1-\beta_2)(\Delta^r)^2, \tag{6}$$

where $\beta_2$ is another decay parameter, $\beta_2 \in [0,1)$, and for **FedAdagrad** is defined as:

$$\boldsymbol{v}^r = \boldsymbol{v}^{r-1} + (\Delta^r)^2. \tag{7}$$

#### 1.5 FedCostWAvg (Mächler et al., 2021):

The aggregation is performed in the same manner as the FedAvg method, but with a modified weighting parameter defined as follows:

$$w_c^r = \alpha \frac{n_c}{N} + (1-\alpha)\frac{k_c^r}{\sum_{i=1}^{C} k_i^r}, \tag{8}$$

where $k_c^r = \frac{\mathcal{L}_c(\boldsymbol{\Theta}_c^{r-1}; \mathcal{D}_c)}{\mathcal{L}_c(\boldsymbol{\Theta}_c^r; \mathcal{D}_c)}$, and $\alpha$ is a hyper-parameter, $\alpha \in [0,1]$. We use $\alpha$=0.5, as suggested in the original study.

1.6 FedPIDAvg (Mächler et al., 2023):

The aggregation is carried out in the same way as the FedAvg algorithm, but with a modified aggregation weight:

$$w_c^r = \alpha \frac{n_c}{N} + \beta \frac{g_c^r}{\sum_{i=1}^{C} g_i^r} + \gamma \frac{h_c^r}{\sum_{i=1}^{C} h_i^r} \tag{9}$$

with $g_c^r = \mathcal{L}_c(\boldsymbol{\Theta}_c^{r-1}; \mathcal{D}_c) - \mathcal{L}_c(\boldsymbol{\Theta}_c^r; \mathcal{D}_c)$, $h_c^r = \sum_{j=0}^{5} \mathcal{L}_c\left(\boldsymbol{\Theta}_c^{r-j}; \mathcal{D}_c\right)$, and $\alpha$, $\beta$, $\gamma$ being hyper-parameters, $\alpha + \beta + \gamma = 1$. The best parameters from the paper were used: $\alpha$=0.45, $\beta$=0.45, $\gamma$=0.1.

1.7 FedBN (Li et al., 2021):

The global model parameters are computed using the FedAvg technique, excluding normalization layers:

$$\boldsymbol{\Theta}_l^{r+1} = \sum_{c=1}^{C} w_c \boldsymbol{\Theta}_{c,l}^r , \tag{10}$$

where $l$ symbolizes non-normalization layers.

1.8 FedMRI:

The aggregation process is executed in the same manner as the FedMean algorithm, but only for the encoder parameters:

$$\boldsymbol{\Theta}_e^{r+1} = \frac{1}{C} \sum_{c=1}^{C} \boldsymbol{\Theta}_{c,e}^r , \tag{11}$$

where $e$ symbolizes encoder. The model used in the study follows the variant referred to as FedMRI† in the original paper by Feng et al., (2023), i.e., the variant without the extra loss component.

1.9 FedBAdam (Feng et al., 2023):

The global model parameters are aggregated using equation (4) of the FedAdam method for all neural network layers, excluding normalization layers:

$$\boldsymbol{\Theta}_l^{r+1} = \boldsymbol{\Theta}_{c,l}^r + \eta \frac{\boldsymbol{m}^r}{\sqrt{\boldsymbol{v}^r} + \tau}, \tag{12}$$

where $l$ symbolizes non-normalization layers, $\boldsymbol{m}^r$ and $\boldsymbol{v}^r$ are calculated according to equation (5) and (6), respectively, but with $\Delta^r = \frac{1}{C} \sum_{c=1}^{C} \left(\boldsymbol{\Theta}_{c,l}^r - \boldsymbol{\Theta}_l^r\right)$.

### 4. Supplementary experimental results

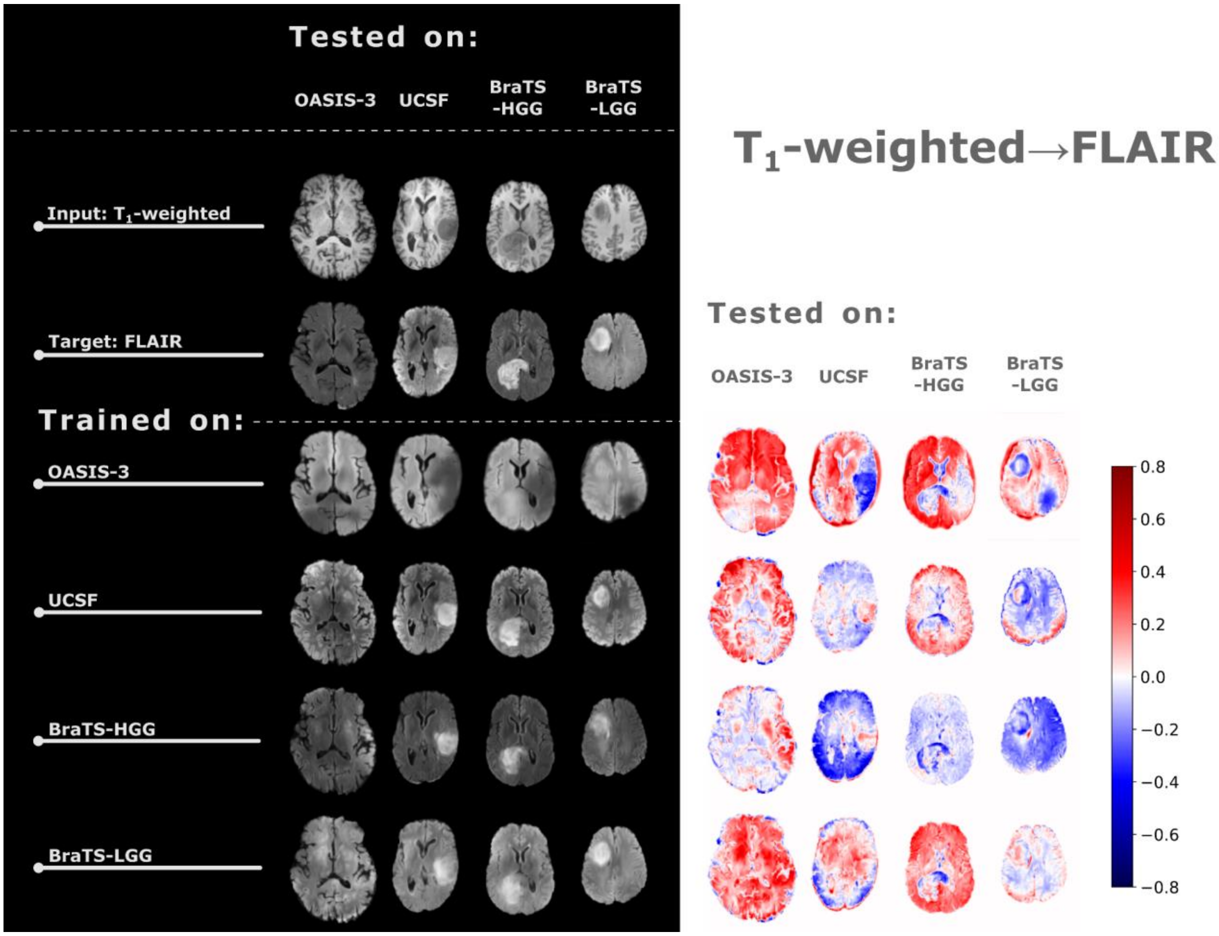


**Supplementary Figure 2a.** Image translation results between T1-weighted and FLAIR MRI in a non-FL way (left panel) and the relative errors calculated between the synthesized data and target images. The first row in the left panel presents the input data, and the second row indicates the target domain (the domain to which the input data is translated). Each subsequent row depicts the results of image translation for the clients (each annotated in the column header) given the training procedure conducted with the data set annotated in the row header.

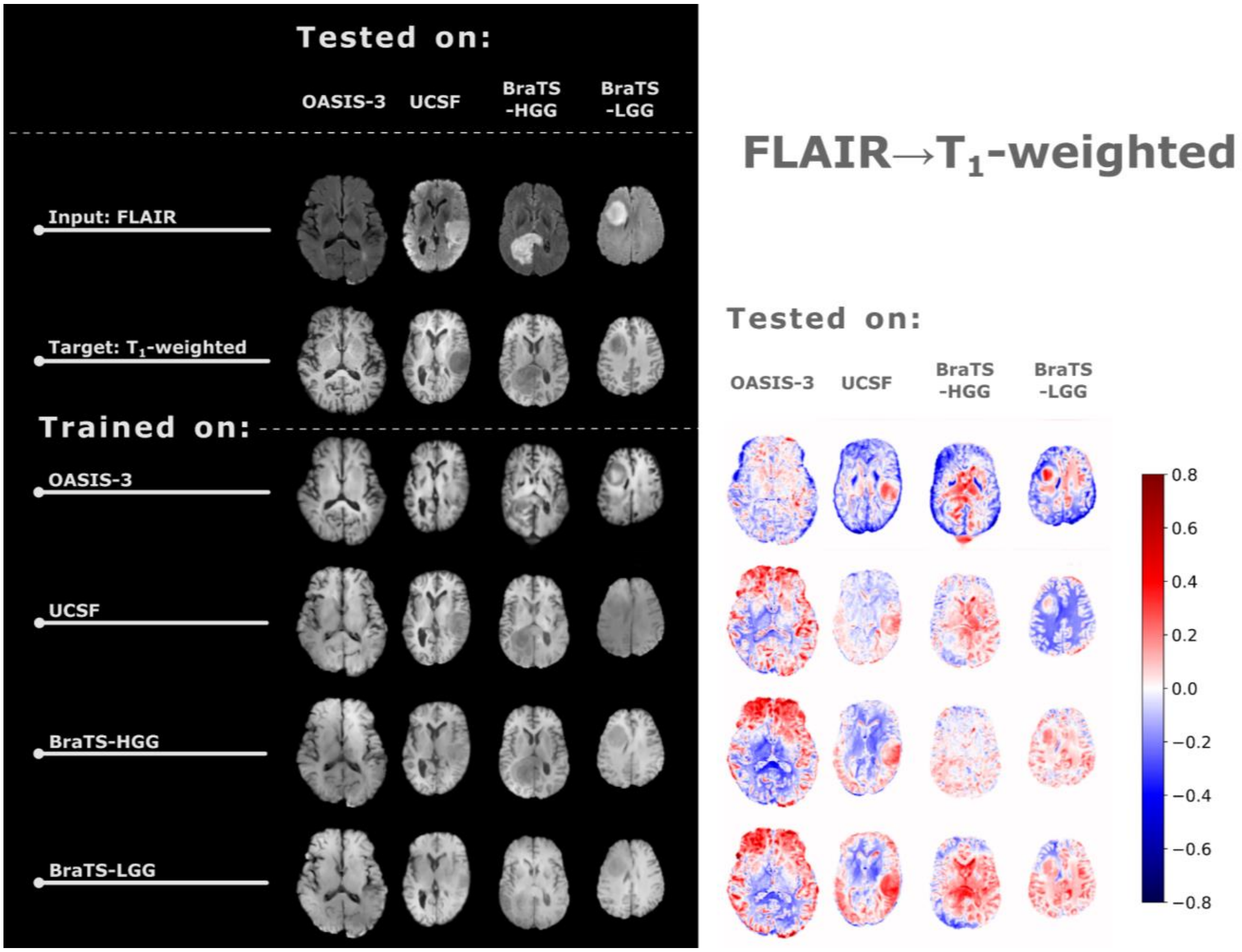


**Supplementary Figure 2b.** (cont.)

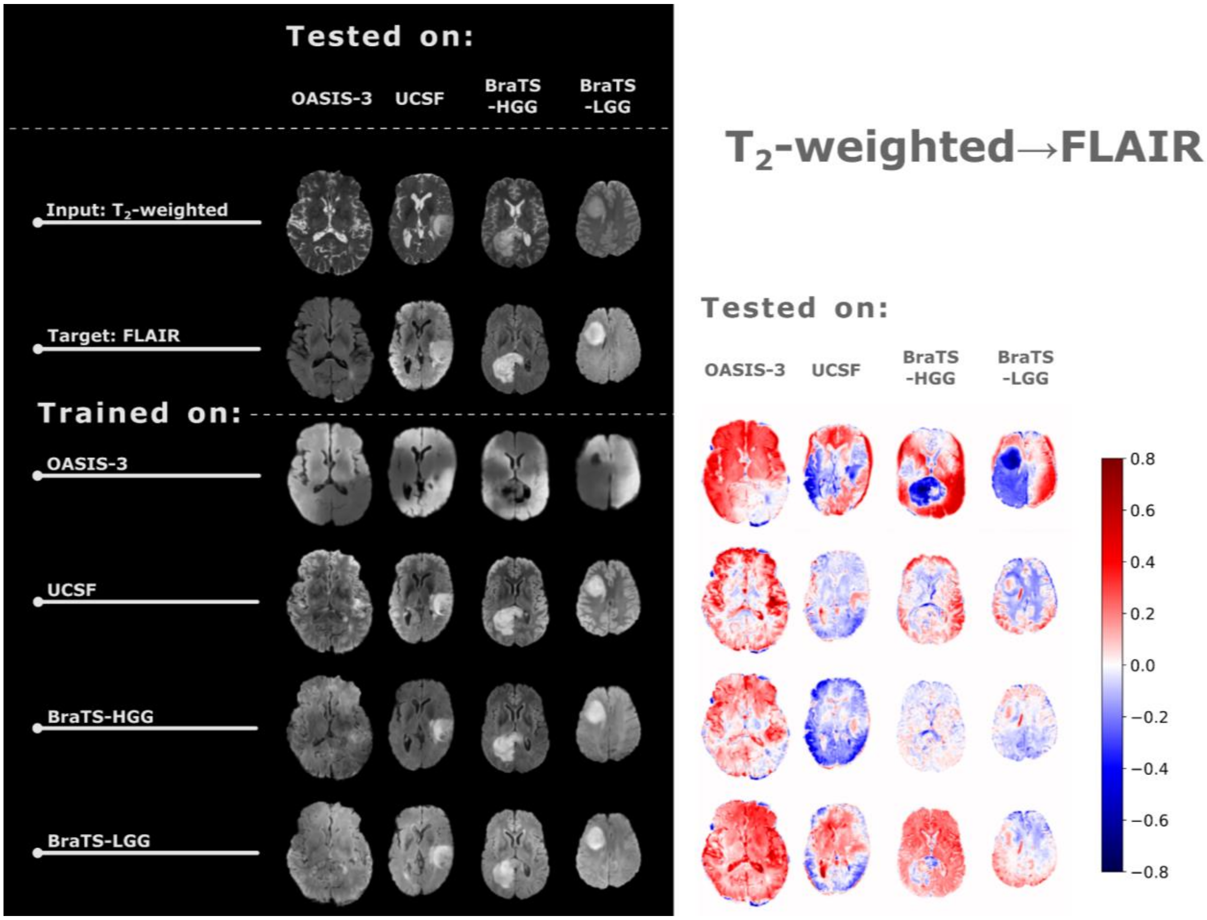


**Supplementary Figure 3a.** Image translation results between T2-weighted and FLAIR MRI in a non-FL way (left panel) and the relative errors calculated between the synthesized data and target images. The first row in the left panel presents the input data, and the second row indicates the target domain (the domain to which the input data is translated). Each subsequent row depicts the results of image translation for the clients (each annotated in the column header) given the training procedure conducted with the data set annotated in the row header.

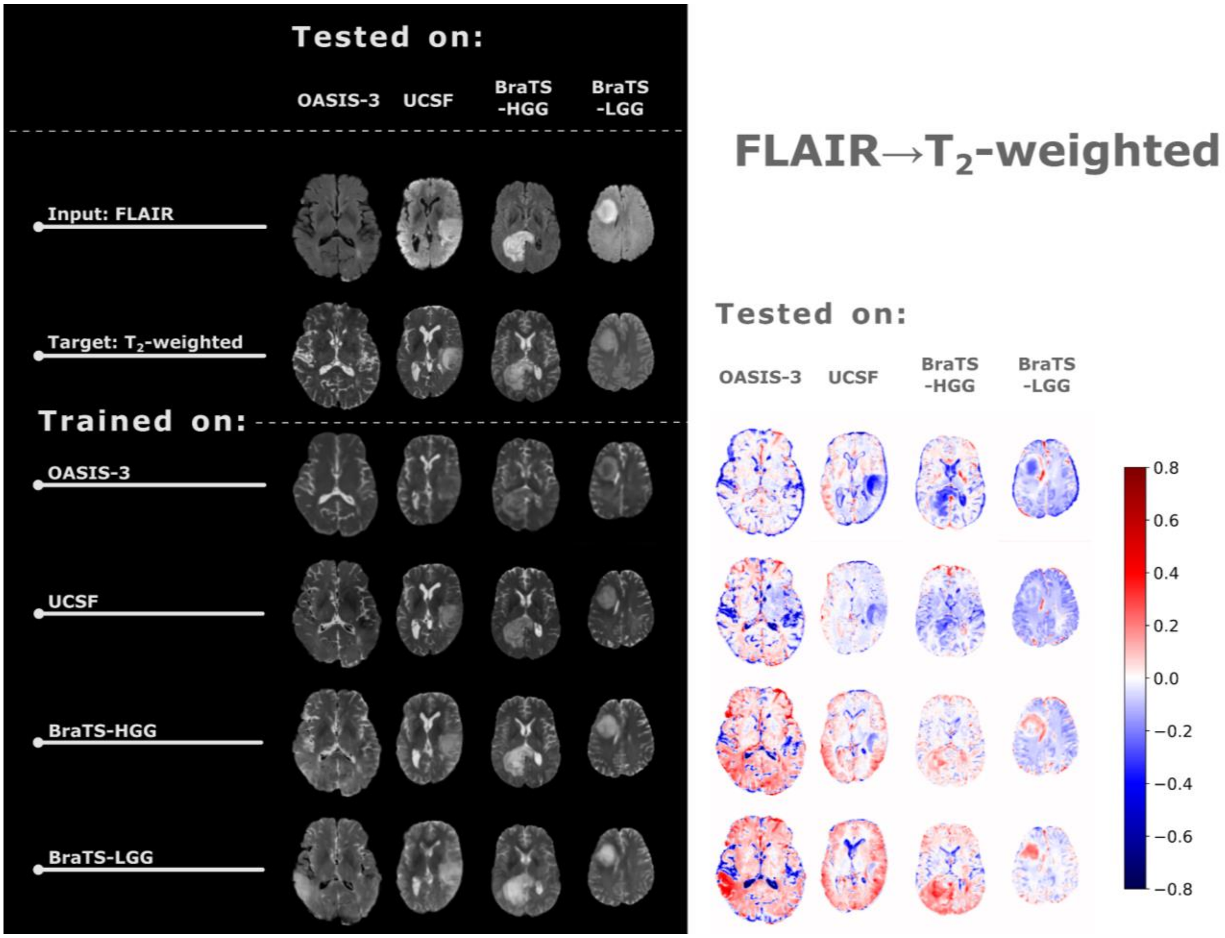


**Supplementary Figure 3b.** (cont.)

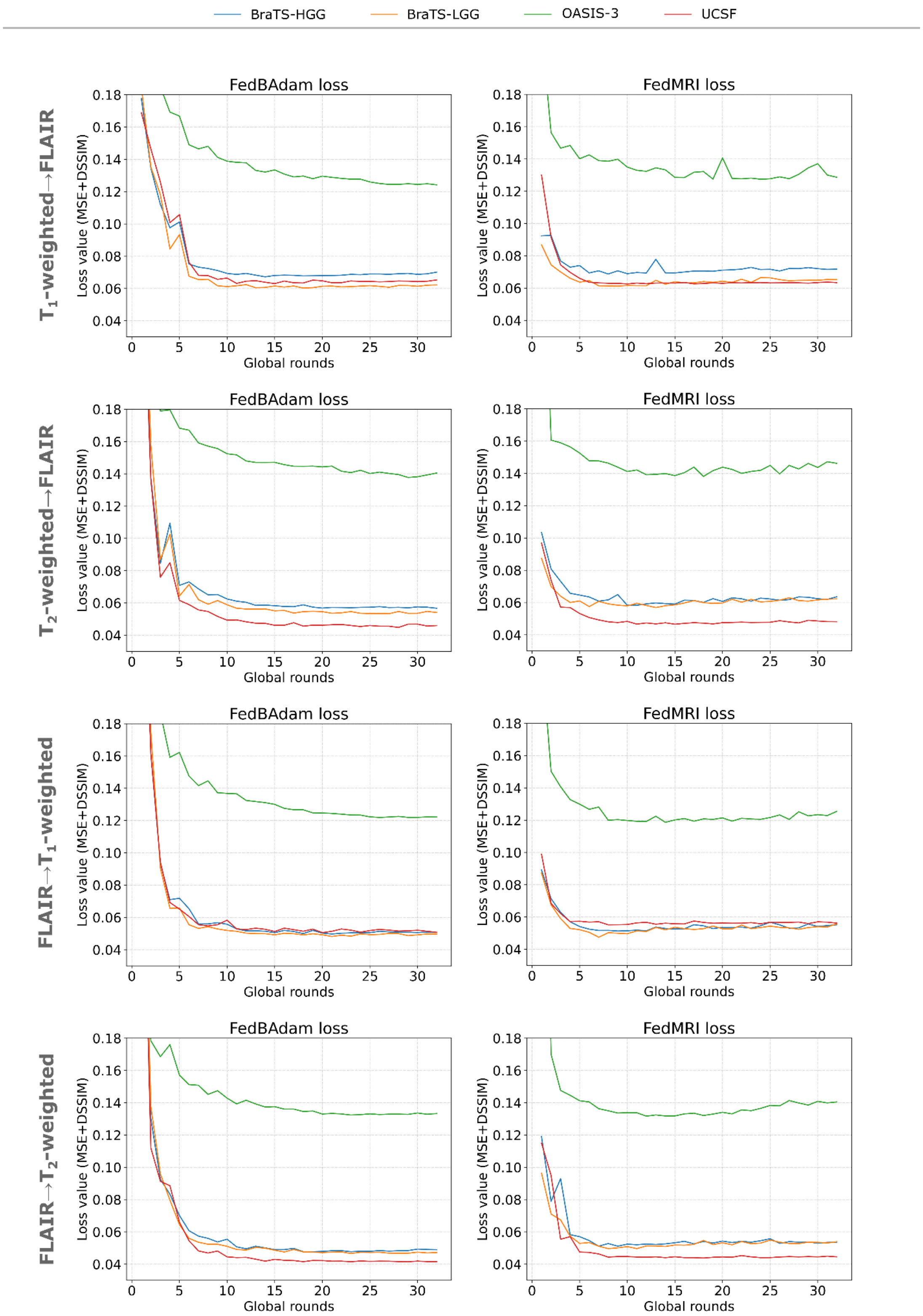


**Supplementary Figure 4.** The change of the loss values as a function of the global round number (global epoch) across the clients involved in the FL-based training for two selected personalized methods (i.e., FedBAdam and FedMRI). The annotations on the left show the direction of image translation.

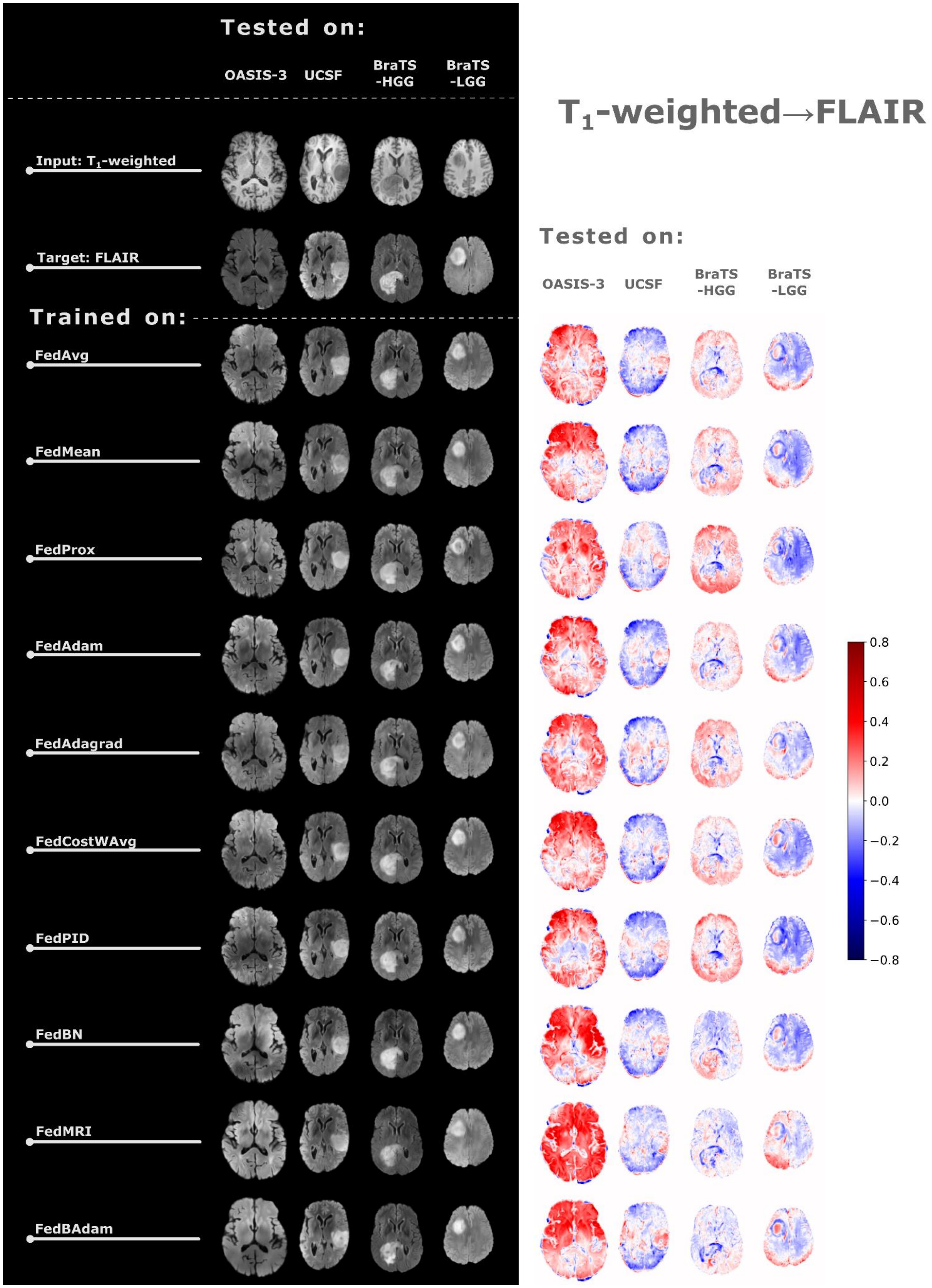


**Supplementary Figure 5a.** Image translation results between T1-weighted and FLAIR MRI in a FL way (left panel) and the relative errors calculated between the synthesized data and target images. The first row in the left panel presents the input data, and the second row indicates the target domain (the domain to which the input data is translated). Each subsequent row depicts the results of the image translation task under the FL-based architecture annotated in the row header for the clients annotated in the column header.

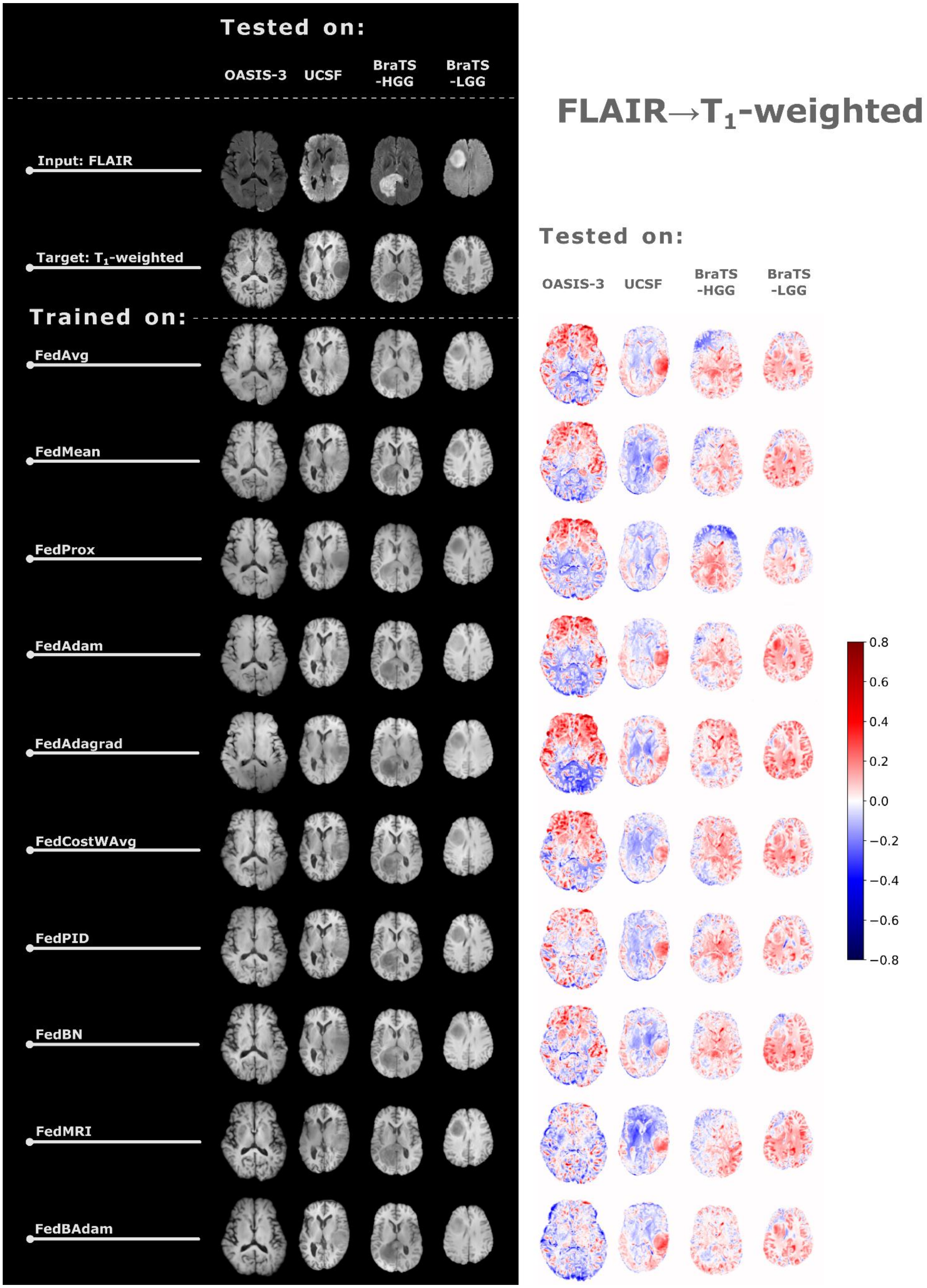


**Supplementary Figure 5b.**

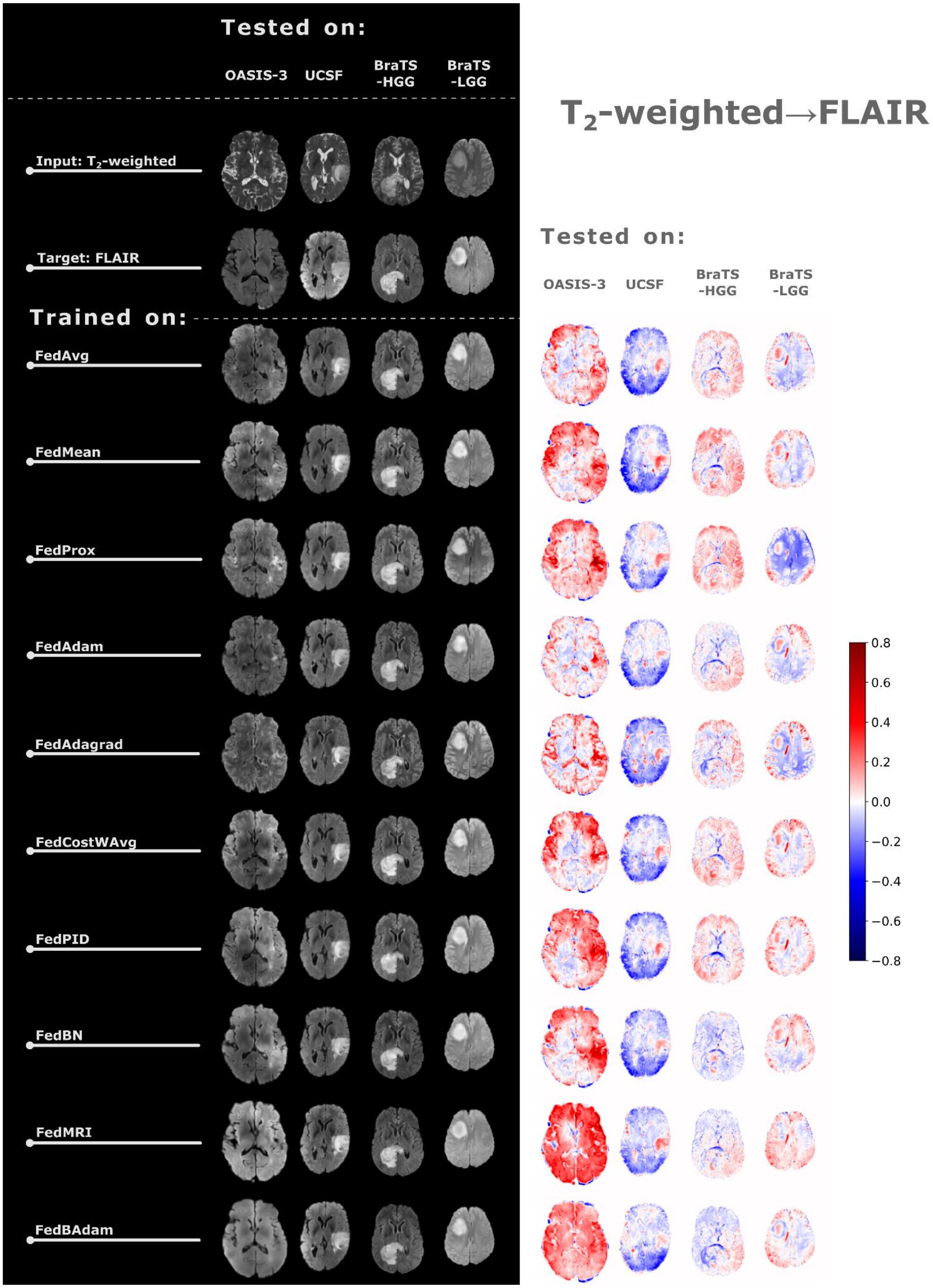


**Supplementary Figure 6a.** Image translation results between T2-weighted and FLAIR MRI in a FL way (left panel) and the relative errors calculated between the synthesized data and target images. The first row in the left panel presents the input data, and the second row indicates the target domain (the domain to which the input data is translated). Each subsequent row depicts the results of the image translation task under the FL-based architecture annotated in the row header for the clients annotated in the column header.

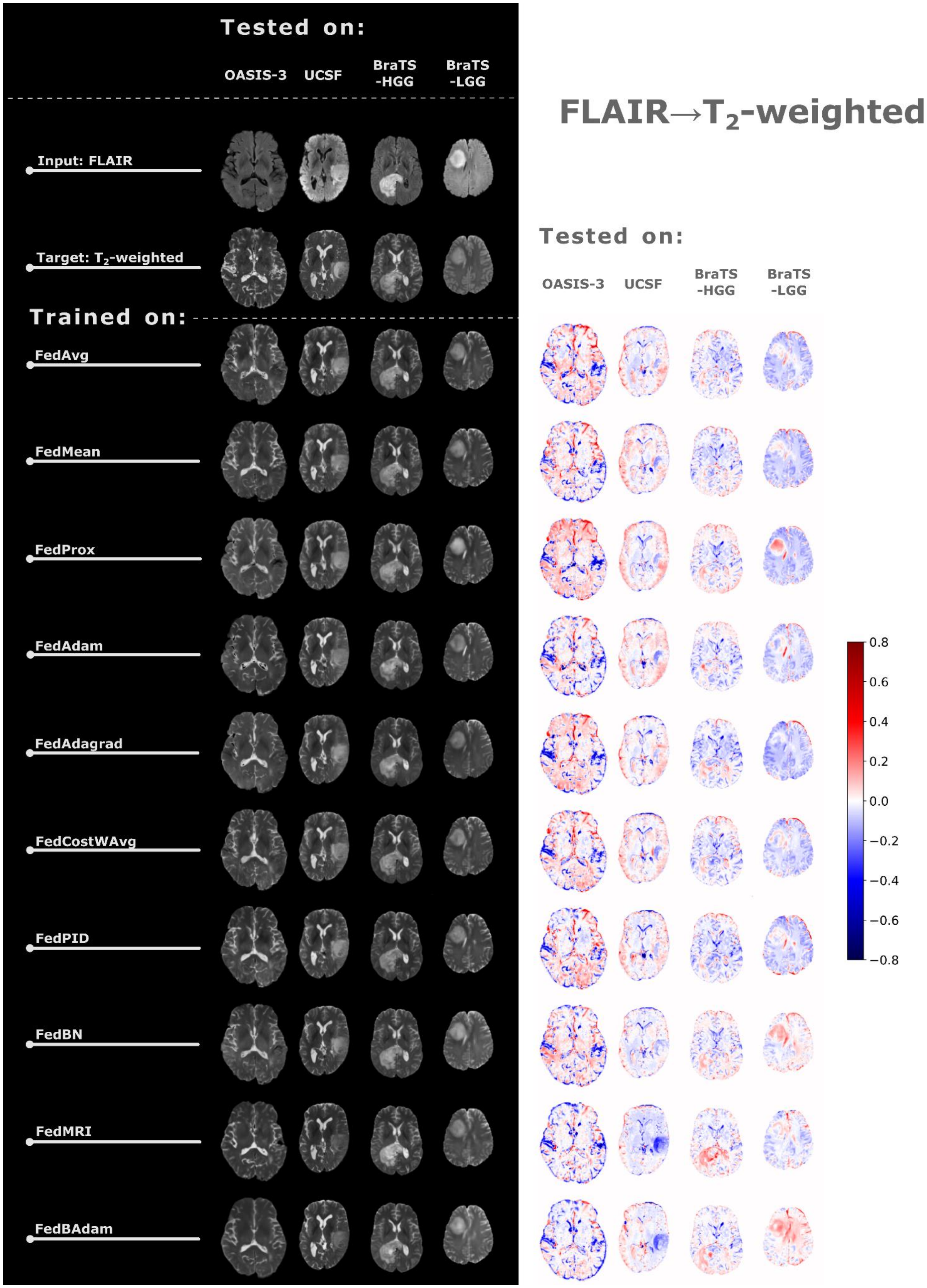


**Supplementary Figure 6b.**

#### $T_1$-weighted → FLAIR

| | OASIS-3 | UCSF | BraTS-HGG | BraTS-LGG | Average |
|---|---|---|---|---|---|
| OASIS-3 | 44.1 ± 27.0 | 50.4 ± 19.1 | 57.6 ± 32.7 | 32.7 ± 12.5 | 46.18 |
| UCSF | 58.7 ± 17.3 | 17.0 ± 9.0 | 35.4 ± 18.0 | 29.6 ± 12.7 | 35.15 |
| BraTS-HGG | 82.0 ± 38.6 | 34.0 ± 15.1 | 26.2 ± 21.6 | 39.8 ± 26.8 | 45.51 |
| BraTS-LGG | 50.8 ± 20.6 | 35.6 ± 14.0 | 44.5 ± 31.1 | 18.5 ± 8.8 | 37.36 |
| FedAvg | 51.5 ± 17.4 | 17.7 ± 8.5 | 22.7 ± 14.9 | 21.0 ± 11.2 | 28.24 |
| FedMean | 40.8 ± 14.6 | 16.5 ± 7.6 | 22.8 ± 15.1 | 18.5 ± 9.3 | 24.65 |
| FedProx | 53.0 ± 18.0 | 19.4 ± 8.8 | 32.2 ± 17.7 | 26.8 ± 14.5 | 32.83 |
| FedAdam | 46.5 ± 15.7 | 19.1 ± 10.0 | 22.3 ± 15.3 | 19.4 ± 9.2 | 26.81 |
| FedAdagrad | 49.4 ± 16.1 | 20.4 ± 7.6 | 25.5 ± 15.6 | 19.3 ± 10.1 | 28.63 |
| FedCostWAvg | 50.6 ± 16.8 | 19.0 ± 8.7 | 23.4 ± 15.2 | 21.7 ± 12.4 | 28.66 |
| FedPID | 41.9 ± 15.8 | 20.1 ± 8.8 | 23.1 ± 15.2 | 18.6 ± 9.6 | 25.91 |
| FedBN | 46.0 ± 22.5 | 16.8 ± 8.1 | 23.7 ± 16.9 | 19.1 ± 10.5 | 26.39 |
| FedMRI | 41.5 ± 31.6 | 16.5 ± 7.6 | 21.5 ± 15.7 | 16.3 ± 7.6 | 23.93 |
| FedBAdam | 31.9 ± 14.5 | 17.6 ± 8.3 | 18.9 ± 12.1 | 15.3 ± 7.4 | 20.94 |

#### $T_2$-weighted→ FLAIR

| | OASIS-3 | UCSF | BraTS-HGG | BraTS-LGG | Average |
|---|---|---|---|---|---|
| OASIS-3 | 56.8 ± 22.6 | 58.2 ± 17.8 | 73.6 ± 25.7 | 66.9 ± 21.5 | 63.87 |
| UCSF | 67.1 ± 18.2 | 15.0 ± 11.9 | 33.4 ± 17.9 | 28.8 ± 12.5 | 36.05 |
| BraTS-HGG | 62.8 ± 21.9 | 20.4 ± 8.6 | 19.3 ± 13.7 | 22.5 ± 13.7 | 31.25 |
| BraTS-LGG | 57.2 ± 22.1 | 35.0 ± 16.1 | 38.5 ± 29.6 | 17.1 ± 9.2 | 36.94 |
| FedAvg | 58.1 ± 19.6 | 14.0 ± 8.0 | 20.8 ± 15.9 | 19.0 ± 10.7 | 28.00 |
| FedMean | 46.5 ± 16.5 | 16.5 ± 8.8 | 23.5 ± 19.6 | 16.4 ± 7.8 | 25.74 |
| FedProx | 59.7 ± 18.2 | 16.6 ± 12.7 | 25.9 ± 18.1 | 26.0 ± 16.8 | 32.02 |
| FedAdam | 51.3 ± 18.7 | 13.0 ± 7.6 | 19.0 ± 15.4 | 17.1 ± 11.1 | 25.11 |
| FedAdagrad | 66.1 ± 22.5 | 17.2 ± 9.6 | 23.3 ± 14.9 | 23.4 ± 13.0 | 32.51 |
| FedCostWAvg | 52.0 ± 16.8 | 15.8 ± 9.4 | 24.0 ± 19.4 | 17.7 ± 8.1 | 27.35 |
| FedPID | 43.9 ± 15.5 | 15.7 ± 7.4 | 23.0 ± 17.7 | 17.1 ± 8.6 | 24.91 |
| FedBN | 51.1 ± 17.3 | 13.6 ± 8.9 | 19.3 ± 13.8 | 16.3 ± 7.9 | 25.08 |
| FedMRI | 42.4 ± 28.1 | 13.9 ± 10.5 | 19.8 ± 13.8 | 17.5 ± 9.6 | 23.41 |
| FedBAdam | 43.2 ± 21.2 | 13.1 ± 8.8 | 17.3 ± 14.5 | 15.1 ± 9.1 | 22.18 |

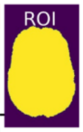


#### FLAIR → $T_1$-weighted

| | OASIS-3 | UCSF | BraTS-HGG | BraTS-LGG | Average |
|---|---|---|---|---|---|
| OASIS-3 | 23.8 ± 12.8 | 46.3 ± 20.5 | 49.7 ± 16.6 | 51.0 ± 18.9 | 42.72 |
| UCSF | 37.3 ± 11.3 | 21.0 ± 16.3 | 25.8 ± 16.8 | 32.2 ± 20.6 | 29.07 |
| BraTS-HGG | 34.4 ± 12.6 | 28.1 ± 26.2 | 17.9 ± 16.1 | 20.1 ± 11.4 | 25.13 |
| BraTS-LGG | 43.4 ± 20.3 | 36.7 ± 33.1 | 30.3 ± 27.2 | 21.4 ± 16.9 | 32.94 |
| FedAvg | 29.0 ± 10.7 | 22.4 ± 21.9 | 17.4 ± 15.8 | 17.4 ± 11.2 | 21.55 |
| FedMean | 25.4 ± 10.8 | 25.9 ± 25.3 | 18.7 ± 15.0 | 18.9 ± 11.4 | 22.24 |
| FedProx | 30.7 ± 15.0 | 25.9 ± 25.7 | 23.3 ± 20.1 | 20.9 ± 13.0 | 25.22 |
| FedAdam | 26.5 ± 9.4 | 23.9 ± 24.5 | 17.3 ± 15.9 | 15.8 ± 10.2 | 20.88 |
| FedAdagrad | 36.6 ± 11.6 | 27.1 ± 25.8 | 17.7 ± 15.4 | 18.3 ± 11.8 | 24.93 |
| FedCostWAvg | 26.5 ± 10.2 | 24.0 ± 21.7 | 17.9 ± 14.2 | 17.5 ± 11.1 | 21.48 |
| FedPID | 22.5 ± 8.8 | 24.8 ± 23.0 | 19.1 ± 14.7 | 20.2 ± 12.6 | 21.63 |
| FedBN | 24.4 ± 10.5 | 22.7 ± 20.1 | 17.5 ± 16.5 | 18.7 ± 12.1 | 20.82 |
| FedMRI | 22.8 ± 9.3 | 22.1 ± 13.8 | 17.2 ± 13.9 | 19.0 ± 12.2 | 20.27 |
| FedBAdam | 20.0 ± 8.6 | 19.2 ± 14.7 | 15.3 ± 14.0 | 16.9 ± 11.5 | 17.84 |

#### FLAIR → $T_2$-weighted

| | OASIS-3 | UCSF | BraTS-HGG | BraTS-LGG | Average |
|---|---|---|---|---|---|
| OASIS-3 | 22.5 ± 9.3 | 22.1 ± 8.4 | 27.7 ± 15.8 | 39.9 ± 25.4 | 28.03 |
| UCSF | 31.1 ± 12.1 | 9.8 ± 6.3 | 22.6 ± 17.6 | 40.1 ± 29.8 | 25.92 |
| BraTS-HGG | 32.5 ± 10.6 | 18.1 ± 8.0 | 12.2 ± 9.7 | 20.2 ± 17.0 | 20.77 |
| BraTS-LGG | 38.7 ± 12.1 | 29.7 ± 12.2 | 15.1 ± 8.5 | 15.8 ± 12.2 | 24.80 |
| FedAvg | 26.4 ± 10.8 | 11.3 ± 6.8 | 12.8 ± 12.4 | 23.3 ± 21.8 | 18.44 |
| FedMean | 22.8 ± 9.5 | 12.0 ± 6.7 | 12.9 ± 12.3 | 23.3 ± 21.3 | 17.77 |
| FedProx | 26.1 ± 10.2 | 13.6 ± 6.8 | 14.2 ± 12.3 | 23.4 ± 21.0 | 19.30 |
| FedAdam | 27.0 ± 10.3 | 11.9 ± 7.0 | 12.2 ± 10.7 | 22.3 ± 21.2 | 18.35 |
| FedAdagrad | 26.1 ± 11.0 | 12.1 ± 7.1 | 14.1 ± 12.6 | 24.8 ± 21.5 | 19.28 |
| FedCostWAvg | 24.6 ± 9.9 | 12.2 ± 7.1 | 11.9 ± 11.3 | 21.8 ± 21.4 | 17.66 |
| FedPID | 23.9 ± 9.9 | 12.9 ± 7.5 | 13.2 ± 11.9 | 22.7 ± 20.6 | 18.19 |
| FedBN | 23.4 ± 10.9 | 9.3 ± 5.3 | 10.9 ± 9.7 | 14.2 ± 11.7 | 14.45 |
| FedMRI | 24.0 ± 11.0 | 10.1 ± 6.3 | 11.6 ± 9.1 | 15.4 ± 11.7 | 15.27 |
| FedBAdam | 20.3 ± 9.2 | 9.2 ± 5.3 | 10.6 ± 8.1 | 12.6 ± 9.2 | 13.17 |

**Supplementary Figure 7.** The MSE (multiplied by a factor of 1000) calculated for synthesized data over the brain region in T1-weighted/T2-weighted→FLAIR translation direction (top tables) and FLAIR →T1-weighted/T2-weighted direction (bottom tables). In all tables the first four rows cover the translation results in a non-FL way, while the remaining rows represent the ten tested FL methods. Each cell represents the MSE ± standard deviation of the error. The last column presents the average computed across all datasets given the training dataset or FL variant.

| | $T_1$-weighted → FLAIR | | | | | $T_2$-weighted→ FLAIR | | | | |
|---|---|---|---|---|---|---|---|---|---|---|
| | OASIS-3 | UCSF | BraTS-HGG | BraTS-LGG | Average | OASIS-3 | UCSF | BraTS-HGG | BraTS-LGG | Average |
| OASIS-3 | 0.44 ± 0.06 | 0.4 ± 0.07 | 0.39 ± 0.07 | 0.44 ± 0.06 | 0.42 | 0.37 ± 0.07 | 0.37 ± 0.05 | 0.36 ± 0.06 | 0.39 ± 0.06 | 0.37 |
| UCSF | 0.3 ± 0.04 | 0.62 ± 0.11 | 0.44 ± 0.07 | 0.47 ± 0.07 | 0.46 | 0.25 ± 0.05 | 0.7 ± 0.1 | 0.48 ± 0.08 | 0.5 ± 0.08 | 0.48 |
| BraTS-HGG | 0.31 ± 0.07 | 0.44 ± 0.07 | 0.53 ± 0.08 | 0.51 ± 0.06 | 0.45 | 0.28 ± 0.06 | 0.55 ± 0.07 | 0.6 ± 0.07 | 0.59 ± 0.07 | 0.51 |
| BraTS-LGG | 0.33 ± 0.05 | 0.47 ± 0.08 | 0.47 ± 0.08 | 0.57 ± 0.07 | 0.46 | 0.29 ± 0.06 | 0.51 ± 0.07 | 0.54 ± 0.09 | 0.62 ± 0.06 | 0.49 |
| FedAvg | 0.36 ± 0.04 | 0.59 ± 0.09 | 0.54 ± 0.08 | 0.57 ± 0.07 | 0.51 | 0.31 ± 0.06 | 0.66 ± 0.08 | 0.59 ± 0.08 | 0.61 ± 0.07 | 0.54 |
| FedMean | 0.4 ± 0.05 | 0.61 ± 0.11 | 0.53 ± 0.07 | 0.58 ± 0.06 | 0.53 | 0.34 ± 0.07 | 0.63 ± 0.07 | 0.58 ± 0.08 | 0.61 ± 0.07 | 0.54 |
| FedProx | 0.37 ± 0.05 | 0.6 ± 0.09 | 0.52 ± 0.08 | 0.57 ± 0.07 | 0.51 | 0.3 ± 0.06 | 0.67 ± 0.09 | 0.59 ± 0.09 | 0.6 ± 0.09 | 0.54 |
| FedAdam | 0.38 ± 0.05 | 0.59 ± 0.1 | 0.54 ± 0.08 | 0.58 ± 0.06 | 0.52 | 0.34 ± 0.07 | 0.69 ± 0.08 | 0.61 ± 0.08 | 0.63 ± 0.07 | 0.57 |
| FedAdagrad | 0.38 ± 0.05 | 0.55 ± 0.07 | 0.53 ± 0.07 | 0.58 ± 0.06 | 0.51 | 0.24 ± 0.06 | 0.63 ± 0.1 | 0.53 ± 0.09 | 0.55 ± 0.1 | 0.49 |
| FedCostWAvg | 0.37 ± 0.05 | 0.56 ± 0.09 | 0.52 ± 0.08 | 0.57 ± 0.06 | 0.51 | 0.33 ± 0.07 | 0.65 ± 0.07 | 0.58 ± 0.08 | 0.61 ± 0.07 | 0.54 |
| FedPID | 0.41 ± 0.05 | 0.56 ± 0.08 | 0.54 ± 0.07 | 0.58 ± 0.06 | 0.52 | 0.35 ± 0.07 | 0.64 ± 0.07 | 0.59 ± 0.08 | 0.62 ± 0.07 | 0.55 |
| FedBN | 0.4 ± 0.05 | 0.61 ± 0.09 | 0.53 ± 0.07 | 0.58 ± 0.06 | 0.53 | 0.35 ± 0.07 | 0.68 ± 0.08 | 0.59 ± 0.07 | 0.62 ± 0.07 | 0.56 |
| FedMRI | 0.44 ± 0.07 | 0.61 ± 0.11 | 0.54 ± 0.08 | 0.57 ± 0.05 | 0.54 | 0.38 ± 0.08 | 0.69 ± 0.09 | 0.57 ± 0.07 | 0.6 ± 0.07 | 0.56 |
| FedBAdam | 0.47 ± 0.06 | 0.6 ± 0.11 | 0.55 ± 0.07 | 0.6 ± 0.06 | 0.56 | 0.39 ± 0.07 | 0.71 ± 0.09 | 0.62 ± 0.07 | 0.65 ± 0.07 | 0.59 |

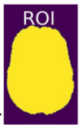


| | FLAIR → $T_1$-weighted | | | | | FLAIR → $T_2$-weighted | | | | |
|---|---|---|---|---|---|---|---|---|---|---|
| | OASIS-3 | UCSF | BraTS-HGG | BraTS-LGG | Average | OASIS-3 | UCSF | BraTS-HGG | BraTS-LGG | Average |
| OASIS-3 | 0.47 ± 0.08 | 0.47 ± 0.08 | 0.44 ± 0.07 | 0.47 ± 0.05 | 0.46 | 0.39 ± 0.09 | 0.52 ± 0.06 | 0.48 ± 0.08 | 0.5 ± 0.08 | 0.47 |
| UCSF | 0.36 ± 0.06 | 0.69 ± 0.09 | 0.58 ± 0.08 | 0.59 ± 0.06 | 0.55 | 0.32 ± 0.08 | 0.73 ± 0.07 | 0.58 ± 0.09 | 0.56 ± 0.09 | 0.55 |
| BraTS-HGG | 0.38 ± 0.07 | 0.62 ± 0.07 | 0.65 ± 0.07 | 0.64 ± 0.05 | 0.57 | 0.31 ± 0.08 | 0.62 ± 0.07 | 0.65 ± 0.08 | 0.64 ± 0.08 | 0.55 |
| BraTS-LGG | 0.36 ± 0.07 | 0.58 ± 0.07 | 0.58 ± 0.09 | 0.66 ± 0.05 | 0.55 | 0.28 ± 0.07 | 0.53 ± 0.08 | 0.59 ± 0.08 | 0.65 ± 0.08 | 0.51 |
| FedAvg | 0.4 ± 0.07 | 0.69 ± 0.09 | 0.66 ± 0.07 | 0.68 ± 0.06 | 0.61 | 0.35 ± 0.09 | 0.71 ± 0.08 | 0.66 ± 0.08 | 0.65 ± 0.08 | 0.59 |
| FedMean | 0.43 ± 0.08 | 0.66 ± 0.09 | 0.63 ± 0.08 | 0.65 ± 0.06 | 0.59 | 0.38 ± 0.1 | 0.68 ± 0.08 | 0.65 ± 0.08 | 0.65 ± 0.08 | 0.59 |
| FedProx | 0.42 ± 0.07 | 0.68 ± 0.07 | 0.64 ± 0.08 | 0.68 ± 0.05 | 0.61 | 0.37 ± 0.09 | 0.7 ± 0.07 | 0.67 ± 0.08 | 0.67 ± 0.09 | 0.60 |
| FedAdam | 0.42 ± 0.08 | 0.7 ± 0.09 | 0.67 ± 0.07 | 0.7 ± 0.06 | 0.62 | 0.36 ± 0.09 | 0.72 ± 0.07 | 0.67 ± 0.08 | 0.67 ± 0.09 | 0.60 |
| FedAdagrad | 0.38 ± 0.06 | 0.67 ± 0.07 | 0.65 ± 0.07 | 0.66 ± 0.06 | 0.59 | 0.35 ± 0.09 | 0.69 ± 0.08 | 0.63 ± 0.08 | 0.64 ± 0.08 | 0.58 |
| FedCostWAvg | 0.42 ± 0.08 | 0.67 ± 0.09 | 0.65 ± 0.07 | 0.67 ± 0.06 | 0.60 | 0.36 ± 0.09 | 0.69 ± 0.08 | 0.66 ± 0.08 | 0.65 ± 0.08 | 0.59 |
| FedPID | 0.43 ± 0.08 | 0.66 ± 0.09 | 0.63 ± 0.07 | 0.64 ± 0.06 | 0.59 | 0.37 ± 0.1 | 0.67 ± 0.08 | 0.63 ± 0.08 | 0.63 ± 0.08 | 0.57 |
| FedBN | 0.44 ± 0.08 | 0.67 ± 0.08 | 0.66 ± 0.08 | 0.67 ± 0.06 | 0.61 | 0.37 ± 0.1 | 0.73 ± 0.07 | 0.67 ± 0.08 | 0.68 ± 0.07 | 0.61 |
| FedMRI | 0.44 ± 0.07 | 0.67 ± 0.08 | 0.64 ± 0.06 | 0.64 ± 0.05 | 0.60 | 0.37 ± 0.1 | 0.72 ± 0.07 | 0.65 ± 0.08 | 0.65 ± 0.07 | 0.60 |
| FedBAdam | 0.47 ± 0.08 | 0.71 ± 0.09 | 0.68 ± 0.07 | 0.7 ± 0.06 | 0.64 | 0.41 ± 0.1 | 0.75 ± 0.07 | 0.68 ± 0.08 | 0.7 ± 0.07 | 0.63 |

**Supplementary Figure 8.** The MSSIM calculated for synthesized data over the brain region in T1-weighted/T2-weighted→FLAIR translation direction (top tables) and FLAIR →T1-weighted/T2-weighted direction (bottom tables). In both tables the first four rows cover the translation results in a non-FL way, while the remaining rows represent the ten tested FL methods. Each cell represents the MSSIM ± standard deviation of the SSIM metric. The last column presents the average computed across all datasets given the training dataset or FL variant. The higher the MSSIM the better.

**$T_1$-weighted → FLAIR**

| | UCSF | BraTS-HGG | BraTS-LGG | Average |
|---|---|---|---|---|
| OASIS-3 | 0.31 ± 0.11 | 0.33 ± 0.1 | 0.32 ± 0.08 | 0.32 |
| UCSF | 0.49 ± 0.15 | 0.43 ± 0.1 | 0.39 ± 0.1 | 0.44 |
| BraTS-HGG | 0.38 ± 0.12 | 0.46 ± 0.09 | 0.43 ± 0.09 | 0.42 |
| BraTS-LGG | 0.38 ± 0.11 | 0.42 ± 0.11 | 0.46 ± 0.1 | 0.42 |
| FedAvg | 0.46 ± 0.13 | 0.48 ± 0.1 | 0.47 ± 0.09 | 0.47 |
| FedMean | 0.46 ± 0.15 | 0.47 ± 0.1 | 0.46 ± 0.09 | 0.47 |
| FedProx | 0.51 ± 0.13 | 0.49 ± 0.11 | 0.47 ± 0.1 | 0.49 |
| FedAdam | 0.45 ± 0.14 | 0.46 ± 0.09 | 0.45 ± 0.09 | 0.45 |
| FedAdagrad | 0.45 ± 0.12 | 0.47 ± 0.1 | 0.47 ± 0.09 | 0.46 |
| FedCostWAvg | 0.44 ± 0.12 | 0.46 ± 0.1 | 0.46 ± 0.09 | 0.45 |
| FedPID | 0.44 ± 0.13 | 0.47 ± 0.1 | 0.48 ± 0.09 | 0.46 |
| FedBN | 0.48 ± 0.14 | 0.46 ± 0.1 | 0.47 ± 0.09 | 0.47 |
| FedMRI | 0.46 ± 0.15 | 0.46 ± 0.1 | 0.47 ± 0.09 | 0.47 |
| FedBAdam | 0.45 ± 0.14 | 0.44 ± 0.1 | 0.46 ± 0.09 | 0.45 |

**$T_2$-weighted → FLAIR**

| | UCSF | BraTS-HGG | BraTS-LGG | Average |
|---|---|---|---|---|
| OASIS-3 | 0.25 ± 0.13 | 0.23 ± 0.13 | 0.24 ± 0.12 | 0.24 |
| UCSF | 0.72 ± 0.12 | 0.56 ± 0.11 | 0.52 ± 0.12 | 0.60 |
| BraTS-HGG | 0.57 ± 0.14 | 0.62 ± 0.1 | 0.55 ± 0.13 | 0.58 |
| BraTS-LGG | 0.5 ± 0.13 | 0.54 ± 0.11 | 0.56 ± 0.11 | 0.54 |
| FedAvg | 0.66 ± 0.12 | 0.61 ± 0.1 | 0.57 ± 0.12 | 0.61 |
| FedMean | 0.65 ± 0.11 | 0.6 ± 0.11 | 0.57 ± 0.11 | 0.61 |
| FedProx | 0.71 ± 0.11 | 0.64 ± 0.11 | 0.58 ± 0.14 | 0.64 |
| FedAdam | 0.7 ± 0.11 | 0.62 ± 0.11 | 0.57 ± 0.13 | 0.63 |
| FedAdagrad | 0.69 ± 0.12 | 0.61 ± 0.1 | 0.54 ± 0.13 | 0.61 |
| FedCostWAvg | 0.67 ± 0.11 | 0.61 ± 0.11 | 0.56 ± 0.12 | 0.61 |
| FedPID | 0.64 ± 0.11 | 0.6 ± 0.12 | 0.57 ± 0.12 | 0.60 |
| FedBN | 0.69 ± 0.11 | 0.61 ± 0.1 | 0.56 ± 0.12 | 0.62 |
| FedMRI | 0.71 ± 0.11 | 0.59 ± 0.11 | 0.55 ± 0.1 | 0.61 |
| FedBAdam | 0.71 ± 0.13 | 0.61 ± 0.11 | 0.59 ± 0.11 | 0.64 |

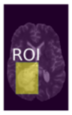


**FLAIR → $T_1$-weighted**

| | UCSF | BraTS-HGG | BraTS-LGG | Average |
|---|---|---|---|---|
| OASIS-3 | 0.38 ± 0.12 | 0.36 ± 0.12 | 0.33 ± 0.1 | 0.36 |
| UCSF | 0.61 ± 0.12 | 0.56 ± 0.11 | 0.53 ± 0.11 | 0.57 |
| BraTS-HGG | 0.55 ± 0.09 | 0.6 ± 0.09 | 0.56 ± 0.11 | 0.57 |
| BraTS-LGG | 0.48 ± 0.1 | 0.53 ± 0.13 | 0.59 ± 0.1 | 0.53 |
| FedAvg | 0.6 ± 0.11 | 0.62 ± 0.09 | 0.6 ± 0.1 | 0.61 |
| FedMean | 0.56 ± 0.11 | 0.58 ± 0.1 | 0.57 ± 0.1 | 0.57 |
| FedProx | 0.62 ± 0.1 | 0.61 ± 0.1 | 0.61 ± 0.09 | 0.61 |
| FedAdam | 0.61 ± 0.11 | 0.63 ± 0.1 | 0.6 ± 0.11 | 0.62 |
| FedAdagrad | 0.59 ± 0.11 | 0.62 ± 0.1 | 0.57 ± 0.1 | 0.59 |
| FedCostWAvg | 0.57 ± 0.11 | 0.6 ± 0.11 | 0.59 ± 0.11 | 0.59 |
| FedPID | 0.57 ± 0.11 | 0.59 ± 0.1 | 0.57 ± 0.1 | 0.57 |
| FedBN | 0.59 ± 0.11 | 0.62 ± 0.09 | 0.6 ± 0.1 | 0.61 |
| FedMRI | 0.58 ± 0.12 | 0.59 ± 0.1 | 0.57 ± 0.09 | 0.58 |
| FedBAdam | 0.62 ± 0.13 | 0.63 ± 0.1 | 0.61 ± 0.1 | 0.62 |

**FLAIR → $T_2$-weighted**

| | UCSF | BraTS-HGG | BraTS-LGG | Average |
|---|---|---|---|---|
| OASIS-3 | 0.53 ± 0.13 | 0.47 ± 0.15 | 0.43 ± 0.13 | 0.48 |
| UCSF | 0.73 ± 0.11 | 0.61 ± 0.1 | 0.52 ± 0.12 | 0.62 |
| BraTS-HGG | 0.6 ± 0.1 | 0.65 ± 0.1 | 0.58 ± 0.11 | 0.61 |
| BraTS-LGG | 0.53 ± 0.11 | 0.6 ± 0.09 | 0.6 ± 0.1 | 0.58 |
| FedAvg | 0.69 ± 0.11 | 0.68 ± 0.09 | 0.62 ± 0.11 | 0.66 |
| FedMean | 0.67 ± 0.1 | 0.66 ± 0.09 | 0.61 ± 0.1 | 0.64 |
| FedProx | 0.67 ± 0.11 | 0.68 ± 0.09 | 0.63 ± 0.11 | 0.66 |
| FedAdam | 0.7 ± 0.11 | 0.68 ± 0.1 | 0.61 ± 0.11 | 0.66 |
| FedAdagrad | 0.69 ± 0.11 | 0.65 ± 0.1 | 0.6 ± 0.11 | 0.65 |
| FedCostWAvg | 0.68 ± 0.11 | 0.67 ± 0.09 | 0.61 ± 0.11 | 0.65 |
| FedPID | 0.66 ± 0.11 | 0.64 ± 0.09 | 0.59 ± 0.11 | 0.63 |
| FedBN | 0.72 ± 0.1 | 0.68 ± 0.09 | 0.64 ± 0.1 | 0.68 |
| FedMRI | 0.71 ± 0.11 | 0.64 ± 0.09 | 0.61 ± 0.09 | 0.65 |
| FedBAdam | 0.73 ± 0.1 | 0.68 ± 0.1 | 0.64 ± 0.09 | 0.68 |

**Supplementary Figure 9.** The MSSIM calculated for synthesized data over the tumour brain region in T1-weighted/T2-weighted→FLAIR translation direction (top tables) and FLAIR →T1-weighted/T2-weighted direction (bottom tables). In both tables the first four rows cover the translation results in a non-FL way, while the remaining rows represent the ten tested FL methods. Each cell represents the MSSIM ± standard deviation of the SSIM metric. The last column presents the average computed across all datasets given the training dataset or FL variant. The higher the MSSIM the better.
